# Genomic architecture of species-specific sex pheromone variation and mate choice

**DOI:** 10.64898/2025.12.11.693631

**Authors:** Weizhao Sun, Marília Fischer, Alexandra Tzipin, Eyal Privman, Jürgen Gadau, Jan Buellesbach

**Affiliations:** Institute for Evolution & Biodiversity, University of Münster, Hüfferstr. 1, DE-48149 Münster, Germany; Department of Evolutionary and Environmental Biology, Institute of Evolution, University of Haifa, Haifa, Israel; Life Science Systems, Technical University of Munich, Hans-Carl-von-Carlowitz-Platz 2, D-85354 Freising, Germany

**Keywords:** comparative genomics, quantitative trait loci, RAD sequencing, gene mapping, sexual signaling, chemical communication, prezygotic isolation, speciation, *Nasonia*, Hymenoptera

## Abstract

Speciation often begins with subtle changes in sexual communication and mate choice, yet the genetic basis of these early reproductive barriers remains largely unresolved. Here, we dissect the chemical and genomic architecture underlying species-specific female cuticular sex pheromones and their role in mate choice in two closely related parasitoid wasp species. Using high-resolution quantitative trait locus (QTL) mapping, fine-scale chemical profiling, and behavioral assays on hybrid populations, we identify three methyl-branched alkanes as key pheromonal compounds that strongly predict male mating preferences. QTL explaining their species-specific variation partially co-localize with genomic regions enriched for candidate genes involved in cuticular hydrocarbon biosynthesis, including cytochrome P450 enzymes and fatty acid synthases. These findings suggest a potential mechanistic link between the variation in these gene clusters and species-specific chemical signaling. Further comparative analyses reveal hitherto unrecognized sex-specific regulatory effects, cytoplasmic influences, and complex genomic interactions shaping the variation in these pheromonal profiles. This study is among the first to connect individual pheromone compounds with mate choice and their underlying genomic basis in a haplodiploid system, offering a rare glimpse into how chemical divergence can reinforce behavioral isolation and potentially drive early speciation.

## Introduction

Understanding how new species arise remains a central question in evolutionary biology, despite more than 150 years since Charles Darwin’s seminal work on speciation through natural selection (Darwin 1859; Shapiro et al. 2016; Shaw et al. 2024). Speciation is widely recognized to involve barriers to interspecific reproduction, which are typically categorized as pre- or postzygotic depending on whether they act before or after zygote formation (Dobzhansky 1937; Coyne and Orr 2004; Merrill et al. 2024). Postzygotic barriers reduce hybrid fitness after fertilization (Orr and Turelli 2001; Frayer and Payseur 2024), whereas prezygotic barriers prevent interspecific mating through ecological, physiological, or behavioral isolation mechanisms (Matute 2014; Uy et al. 2018). Among these, behavioral isolation, which is often driven by divergence in sexual communication, is considered to play a key role in early speciation events (Ortiz-Barrientos et al. 2009; Merrill et al. 2024). Chemical signaling, the most ancient and widespread form of communication, can thereby play a pivotal role in maintaining species-specific communication channels (Johansson and Jones 2007; Wyatt 2014). Particularly insects have exploited chemical signaling as their primary mode of communication (Greenfield 2002; Buellesbach et al. 2018a). However, despite their ecological and evolutionary relevance, the exact chemical and genetic mechanisms governing reproductive isolation remain largely elusive in most insect taxa (Bradbury and Vehrencamp 2011; Servedio and Boughman 2017).

The parasitoid wasp genus *Nasonia* provides a powerful model for studying these processes due to its genetic tractability and well-characterized ecology (Werren et al. 2010; Lynch 2015; Buellesbach et al. 2017). In this species complex, well-studied postzygotic isolation contrasts with far-less understood prezygotic isolation (Gadau et al. 1999; Gibson et al. 2010; Buellesbach et al. 2014). The four naturally occurring *Nasonia* species are all infected with intracellular *Wolbachia* bacteria, leading to bidirectional cytoplasmic incompatibility in most interspecific crosses that result in a substantial to total reduction of hybrid offspring (Breeuwer and Werren 1990; Gadau et al. 1999; Bordenstein et al. 2001). However, their *Wolbachia* infections can be antibiotically cured, rendering hybrids between all four species obtainable under laboratory conditions (Richardson et al. 1987; Bordenstein and Werren 1998). This already indicates that prezygotic reproductive isolation between the different *Nasonia* species is incomplete. But in natural populations, where *Wolbachia* persists, the complete absence of prezygotic isolation would impose severe fitness costs (Bordenstein and Werren 2007; Raychoudhury et al. 2010). Preference of con- over heterospecific mating partners would be a likely consequence to counteract these costs, favoring mechanisms for rapid species recognition and assortative mating (Rundle et al. 2005; Smadja and Butlin 2009; Matute 2014).

*Nasonia* wasps offer a unique opportunity to study species and mate recognition mechanisms due to their diverse and well-studied chemical communication systems (Niehuis et al. 2013; Buellesbach 2018; Mair and Ruther 2019). Particularly female cuticular hydrocarbons (CHCs), complex blends of long-chained lipids coating the insect cuticle, have emerged as key sexual signals triggering courtship and copulation in conspecific males (Steiner et al. 2006; Buellesbach et al. 2013; Sun et al. 2023). However, exactly which CHC components contribute to mate discrimination and therefore prezygotic reproductive isolation could not be resolved so far (Mair et al. 2017; Buellesbach et al. 2018b). Furthermore, the genetic architecture underlying *Nasonia* CHC variation has been mostly studied in males, exploiting their haploidy, but preventing a direct causal link to female CHC profiles and their function in species recognition and mate preference (Niehuis et al. 2011; Buellesbach et al. 2022). Nevertheless, some recent progress has been made in the characterization of candidate genes influencing female CHC profiles and the encoded sexual attractiveness in single *Nasonia* species (Wang et al. 2022; Sun et al. 2023; Sun et al. 2025). However, a comprehensive understanding of the genetic basis of species-specific female CHC variation and its potential impact on prezygotic isolation and mate choice has remained largely elusive so far (Wang et al. 2015; Buellesbach et al. 2022). This would however be instrumental for uncovering and assessing how subtle chemical divergence can reinforce species boundaries and drive the early stages of speciation (Buellesbach and Schmitt 2015; Liu et al. 2023).

In this study, we address this knowledge gap by focusing on the two *Nasonia* species *N. longicornis* and *N. giraulti*, which differ markedly in female CHC profile variation and male mate discrimination behavior. While *N. longicornis* males can reliably distinguish between con- and heterospecific female CHC profiles (see also Fig. 1C), *N. giraulti* males do not exhibit such discrimination behavior (Buellesbach et al. 2013; Giesbers et al. 2013; Mair et al. 2017). Exploiting their haplo-diploid sex determination and interfertility after removing their *Wolbachia* infections, we generated F_2_ hybrid males and backcrossed them to the parental strains to establish clonal female sibship (CFS) lines. These genetically and phenotypically mosaic females were subjected to an integrative analysis combining high-resolution quantitative trait locus (QTL) mapping, chemical profiling and male mate choice assays. This comprehensive approach enabled the identification of individual pheromonal CHC compounds genetically mapped and functionally linked to species-specific male mate preference. We thus uncovered several genomic hotspots enriched with candidate genes for the potential biosynthesis of these pheromonal compounds. Furthermore, through fully leveraging the haplo-diploidy of our study species, we reveal previously unknown effects of cyto-nuclear interactions and sex-specific regulation on the analyzed chemical profiles in their entirety. Our findings provide rare empirical evidence for the contribution of subtle divergences in sexual signaling to mate choice, behavioral isolation and, potentially, the early stages of speciation.

**Figure 1.**
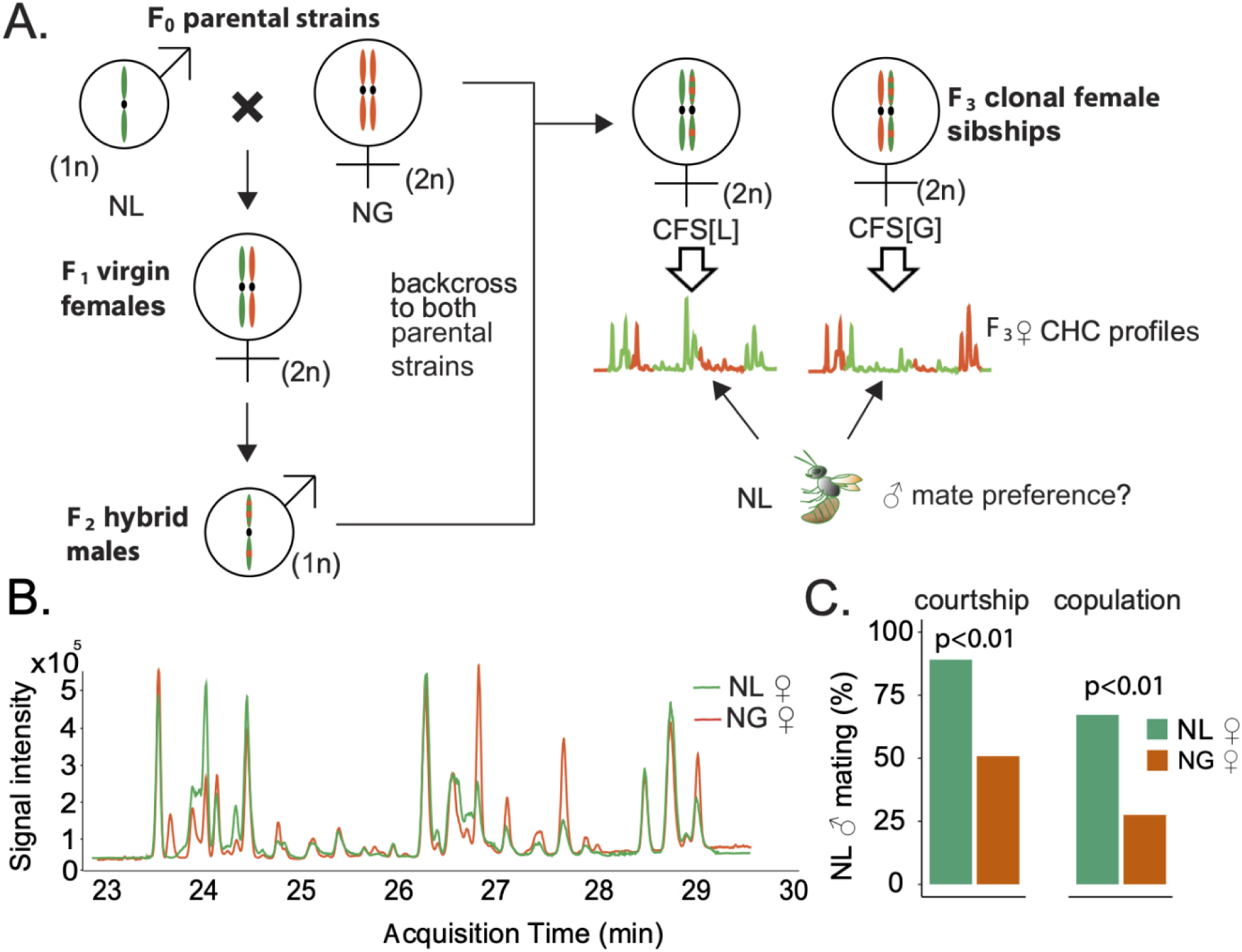
Hybrid crossing scheme, species-specific CHC variation and male mating preference between *Nasonia longicornis* (NL) and N. giraulti (NG). A.) Crossing scheme used to generate clonal female sibship (CFS) lines and schematics for our experimental set-up (for details see Materials and Methods). NL is represented in green, NG in orange, color codes are consistent across panels. B.) Representative female Flame Ionization Detection (FID) chromatograms highlighting quantitative cuticular hydrocarbon (CHC) differences between NL and NG. C.) NL males display significantly higher courtship and copulation rates on NL (N=119) than on NG females (N=120). Significant differences were assessed with Fisher’s exact tests.

## Results

### 1. Generation of clonal female sibships

We generated clonal female sibship (CFS) lines by first crossing interfertile (*i.e., Wolbachia*-free) laboratory strains of *Nasonia longicornis* (NL) and *N. giraulti* (NG). The resulting F_1_ hybrid females were constrained to lay haploid eggs, producing recombinant F_2_ hybrid males that were individually backcrossed to females from both parental strains (Fig. 1A).

This crossing scheme yielded 126 F_2_ hybrid males in total, each fathering at least 20 clonal daughters per CFS line, enabling the assessment of their respective phenotypic variation through genotyping their haploid fathers (Fig. 1B; Fig. S1A; Table S1). The deliberate use of highly inbred parental strains ensured that genetic variation in the generated CFS lines originated almost exclusively from their recombinant F2 fathers (Gadau et al. 2008; Werren and Loehlin 2009). To account for any potential additional genetic variation not exclusively explained through the respective haplotypes, we tracked the respective parental genetic and cytoplasmic background in each CFS for all subsequent analyses (CFS[L] and CFS[G], compare to Fig. 1A).

Backcross success differed markedly between parental backgrounds: 85.4% of NG females mated with hybrid F_2_ males produced female offspring for CFS lines, compared to only 41.7% of NL females (Fig. S2A). NL females also produced a significantly higher proportion of diapause larvae (61.7%) than NG females (10.7%) (Fig. S2B). These types of larvae do not develop into adults but enter a prolonged developmental arrest stage (Schneiderman and Horwitz 1958; Saunders 1965; Foley et al. 2025). Consequently, CFS lines derived from NL females contained fewer individuals than those from NG females (Fig. S2D), indicating stronger and asymmetric postzygotic isolation skewed towards NL (Breeuwer and Werren 1995; Ellison et al. 2008; Gibson et al. 2013).

### 2. Genetic architecture underlying species-specific CHC variation

Genotyping of 332 SNP markers from the 126 F_2_ hybrid males mapped to five linkage groups spanning 787.4 cM (Fig. S3). High-resolution QTL mapping revealed 88 loci associated with species-specific variation in female CHC profiles: 41 in CFS[G] and 47 in CFS[L], collectively explaining variation in 51 of 54 detected CHC compounds (Fig. 2A, Table 1). Most compounds were associated with a single QTL, though some exhibited two loci, and one compound (11,x-DiMeC33) mapped to three distinct QTL in CFS[G] (Fig. 2C). Several genomic regions exhibited high QTL density, particularly on chromosome 1, suggesting hotspots for either clusters of biosynthetic genes or highly pleiotropic loci governing CHC variation (Fig. 2B; Table 1; Buellesbach et al. 2022). On the other chromosomes, however, the distributions of individual QTL partially differed considerably between the two female mapping populations (Fig. 2A-B). Only 14 QTL are shared between CFS[G] and CFS[L], which notably all govern the variation in alkanes with one to three methyl-branches (Fig. 2A and Table 1). This leaves 27 unique QTL in the CFS[G] and 33 in the CFS[L] lines (Fig. 2D), with the vast majority in the latter co-localizing at a genomic hotspot on chromosome 3 (Fig. 2A-B).

**Figure 2.**
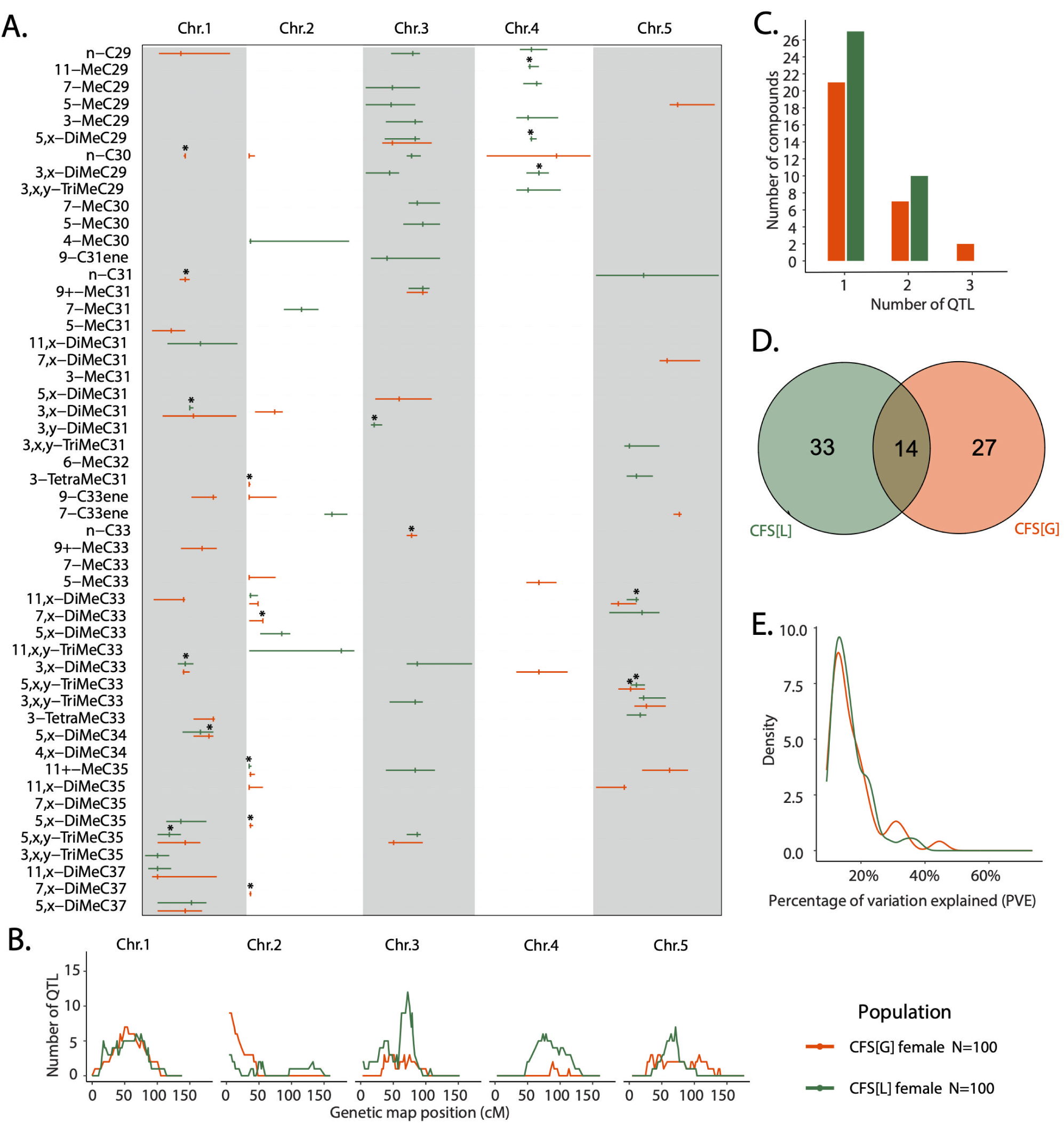
Genomic architecture underlying species-specific CHC variation. A.) QTL positions associated with CHC abundance variation. Bars indicate 95% Bayesian credible intervals, tick marks label the genetic marker with the highest LOD (logarithm of the odds) score within each QTL region. LOD scores higher than 4.85 (explaining >20% phenotypic variation) are indicated by astericks. QTL identified in clonal female sibship (CFS) lines with an NL cytoplasmic background (CFS[L]) are represented in green, the ones with an NG cytoplasmic background are colored in orange (CFS[G]), respectively, color codes are consistent across panels. B.) Number of QTL identified across the genetic map for each mapping population. C.) Distribution of CHC compounds for which 1, 2 or 3 QTL were identified across mapping populations. D.) Venn diagram of unique and shared QTL among the two mapping populations. QTL were considered as shared when their Bayesian credible intervals overlapped. E.) Percentage of variation explained (PVE) by the identidied QTL (N=100 for CFS[L] and CFS[G]).

**Table 1.**
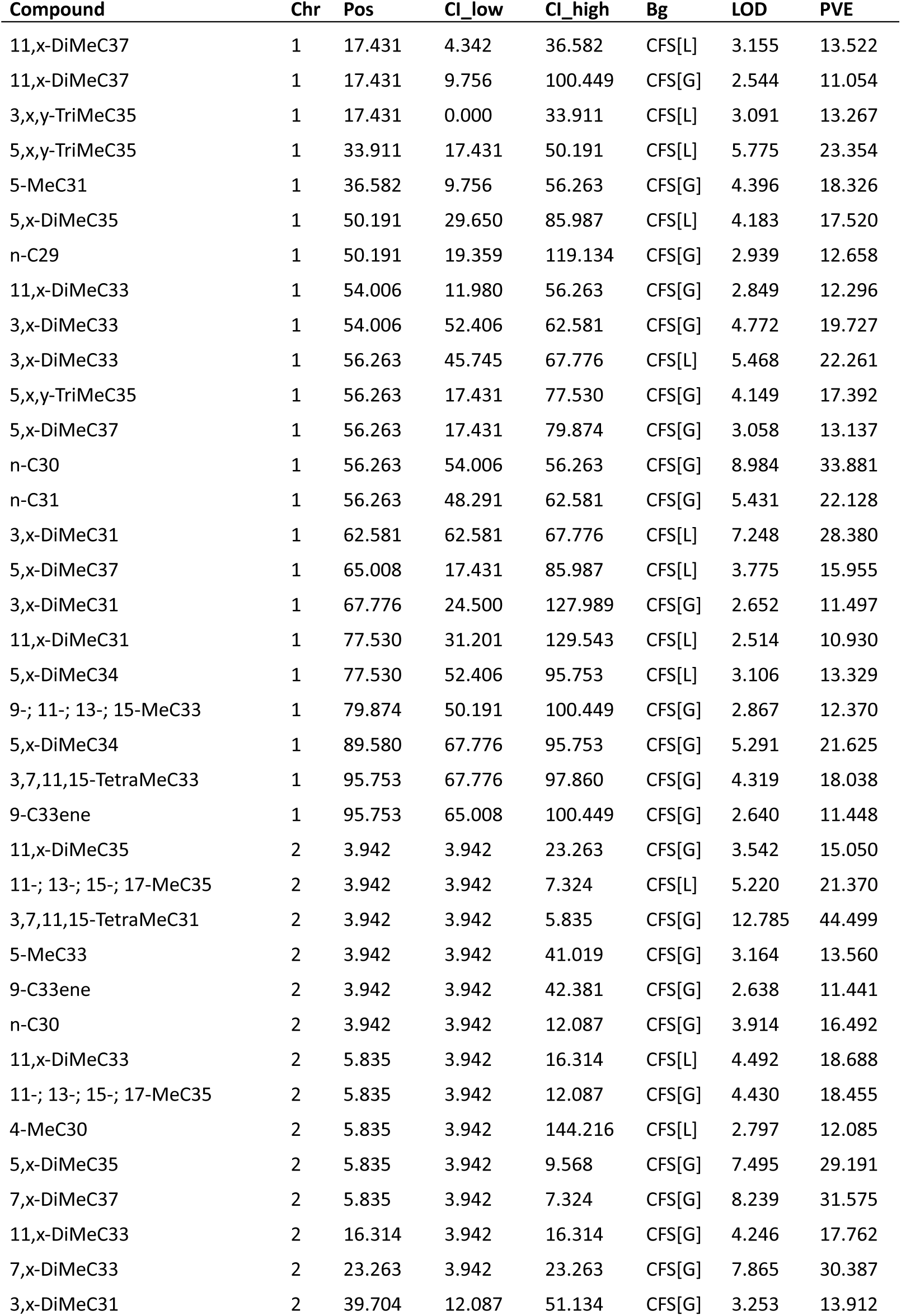
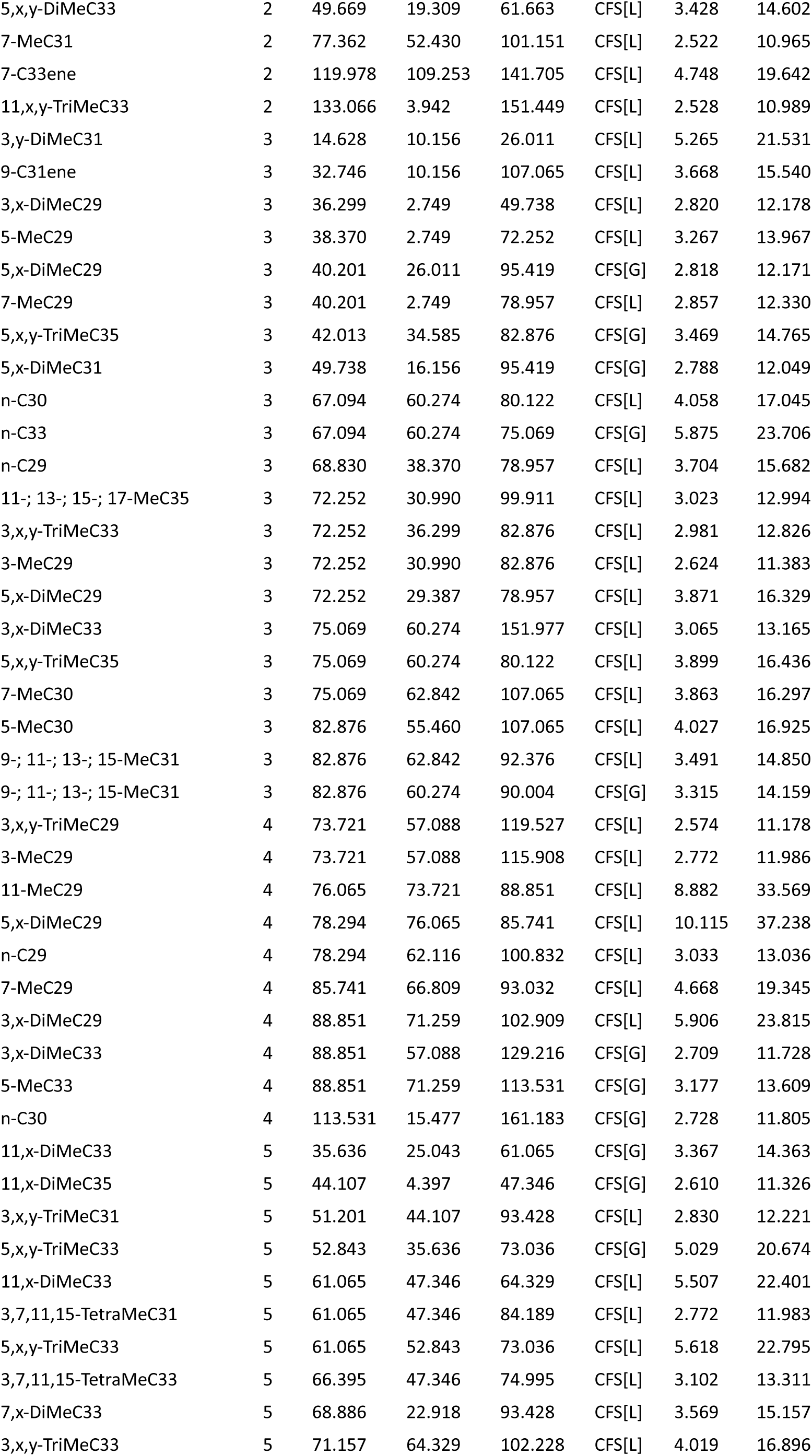
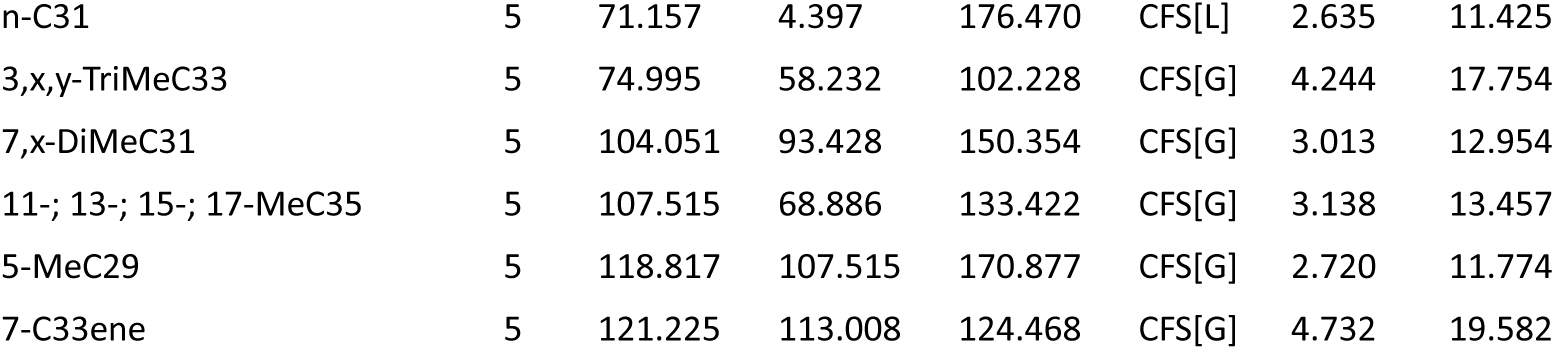
Quantitative trait loci (QTL) associated with species-specific CHC variation. This table lists the total of 171 QTL identified across clonal female sibship (CFS) mapping populations of both genetic backgrounds (NG: [G]; NL: [L]). For each QTL, the following information is provided: chromosome (Chr), peak marker position (Pos) in centimorgan (cM), lower and upper bounds of the 95% Bayesian credible interval (CI_low and CI_high, respectively), mapping population background (CFS[G]; CFS[L]), logarithm of the odds score (LOD), and percentage of variance explained (PVE).

Percentage of variation explained (PVE) by individual QTL ranged primarily between 10–25%, though several loci exhibited higher PVE, indicating large-effect QTL (Fig. 2E). Several of these large effect QTLs overlapped, revealing particularly promising genomic hotspots governing biosynthesis and variation of structurally similar CHCs. For instance, QTLs for 7,x-DiMeC33, 7,x-DiMeC35 and 7,x-DiMeC37, only differing in their respective chain length, co-localized on chromosome 2 in the CFS[G] mapping population, suggesting pleiotropic genes or gene clusters influencing chain-length variation.

### 3. Key CHC compounds correlating with male mate preference

CHC profiles of parental NL and NG females were clearly distinct from both CFS lines, which clustered intermediately but closer to their respective parental female strain from which their cytoplasmic background derived (Fig. 3A, see also Fig. 1A). Behavioral assays revealed that the discriminatory males from the NL strain exhibited higher courtship and copulation frequencies toward CFS[G] than toward CFS[L] females (Fig. 3B).

**Figure 3.**
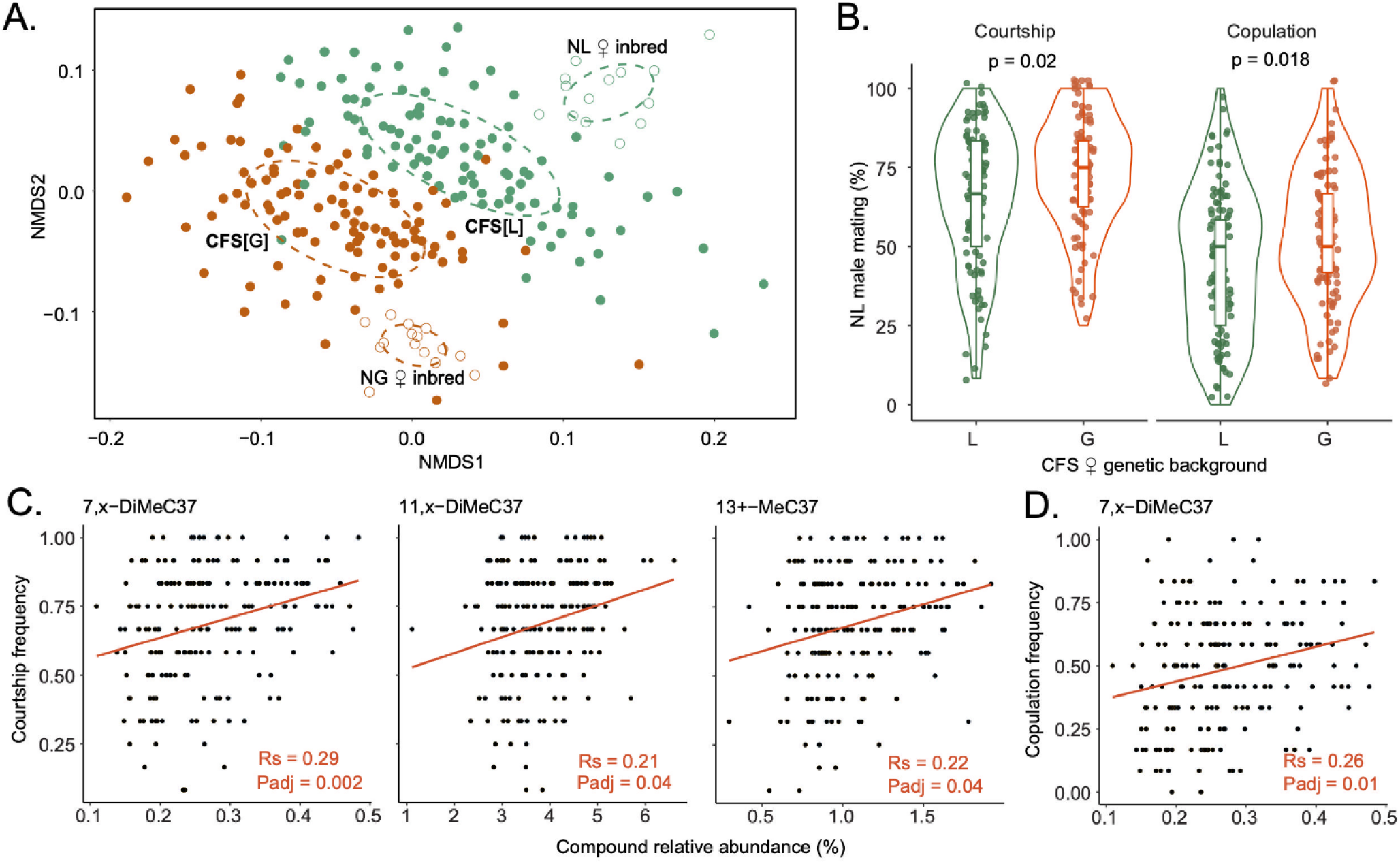
Key CHC compounds correlated with NL male mate preference. A.) Non-Metric Multidimensional Scaling (NMDS) plot demonstrating the divergence of CHC profiles of NL (green open circles, N=10), NG (orange open circles, N=10), CFS[L] (green filled circles, N=101) and CFS[G] (orange filled circles, N=99) females. Each data point for NL and NG females represents the CHC profile of a single individual, whereas each data point for the CFS[L] and CFS[G] populations represents the average CHC abundances of 6-12 individuals. Bray–Curtis dissimilarity model, stress value = 0.23. B.) Violin plots summarizing NL male mating frequencies on CFS[L] (green, N=101) and CFS[G] (orange, N=99) populations. Each point represents the average mating frequency of NL males on 12 individual females from a single respective CFS population. Significant differences (p<0.05) were assessed with Benjamini-Hochberg corrected Mann-Whitney U tests. C-D.) Significant Spearman rank correlations between relative abundances of individual CHC compounds in CFS populations and NL courtship/copulation frequencies (Benjamini-Hochberg corrected for 54 CHC compounds).

To pinpoint individual pheromonal CHC compounds that contribute to species-specific male preference, we correlated their relative abundances in CFS females with NL male courtship and copulation frequencies (Table S2). The abundances of three methyl-branched compounds with identical chain length, *i.e.,* 11,x-DiMeC37 (p<0.05), 13^+^-MeC37 (p<0.05) and 7,x-DiMeC37 (p<0.01), exhibited significant positive correlations with NL male courtship frequencies (Fig. 3C). Among these, 7,x-DiMeC37 was also significantly correlated with NL male copulation frequency (p<0.05, Fig. 3D).

### 4. Phenotypic correlation networks of the identified pheromonal compounds

Since CHCs are synthesized through additive metabolic pathways involving sequential elongation and double bond or methyl group insertions, we expected correlations among compounds sharing the same carbon chain length and/or CHC compound class, *e.g.* number and position of methyl groups (Blomquist and Bagnères 2010; Holze et al. 2021). We therefore investigated phenotypic correlation networks focusing on our identified key pheromonal compounds in the CFS[G] and CFS[L] female populations. For comparison, we also included a phenotypic CHC correlation network based on their F_2_ hybrid male fathers (Fig. 4).

**Figure 4.**
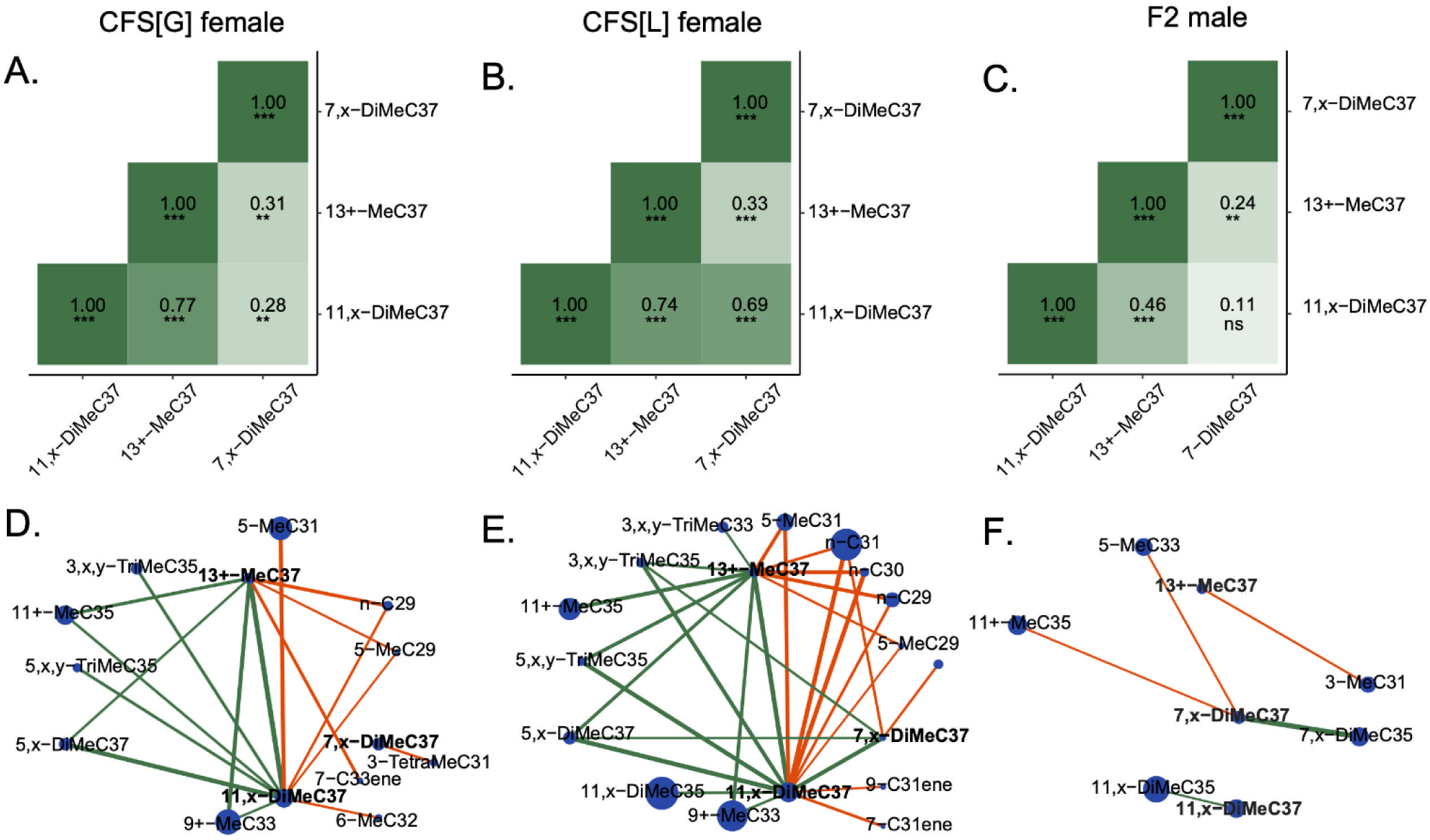
Phenotypic correlation networks of key pheromonal compounds. A-C) Matrices indicating correlations between the three identified key pheromonal compounds across CFS[G], CFS[L] and F_2_ male hybrid populations Green saturation degree reflects Spearman correlation strength in each tile. Significance corrected for multiple testing (Benjamini-Hochberg) and indicated as follows: ns = not significant; * = p<0.05; ** = p<0.01; *** = p<0.001. D-E) Correlation networks of the three focal pheromonal compounds and other CHCs. Node size indicates the relative abundance of individual compounds and edge width indicates of correlation strength, positive correlations are indicated in green, negative ones in red. Only connections with correlation coefficients above 0.5 (positive) and below -0.5 (negative) are displayed in the plot.

Notably, abundances of 7,x-DiMeC37 and 11,x-DiMeC37 were significantly correlated in CFS[G] and CFS[L] females (Fig. 4A-B) but not at all in F_2_ males (Fig. 4C). Furthermore, while the abundance of 7,x-DiMeC37 was similarly correlated with 13^+^-MeC37 in all three populations, 11,x-DiMeC37 and 13^+^-MeC37 abundances were strongly correlated in both CFS populations, but only weakly in F2 males (Fig. 4A-C).

Broader connectivity analyses of the three key pheromonal compounds to the overall identified CHCs showed 17 and 29 linked compounds in CFS[G] and CFS[L], respectively, versus only five in F_2_ males (Fig. 4D–F). The five F2 male correlations were entirely absent from both CFS networks. In contrast, CFS[G] and CFS[L] networks shared several similarities, particularly regarding CHCs connected to 11,x-DiMeC37 and 13^+^-MeC37. In both populations, these compounds showed positive correlations with multiple long-chained CHCs, including 9+-MeC33, 11+-MeC35, 3,x,y-TriMeC35, 5,x,y-TriMeC35, and 5,x-DiMeC37.

Conversely, both networks displayed negative correlations between these two compounds and shorter-chained CHCs (*e.g., n*-C29, 5-MeC29, 5-MeC31), with additional negative correlations with compounds such as *n*-C30, *n*-C31, 7-C31ene and 9-C31ene in the CFS[L] female network. However, network structure around 7,x-DiMeC37 differed substantially between CFS backgrounds, indicating divergent regulatory interactions.

### 5. Co-localization between QTL for male mate preference and key pheromonal compounds

We detected a single QTL for NL male courtship frequency exclusively in CFS[L], located on chromosome 1 (13.3 to 138.2 cM, Fig. 5A). To further explore the behavioral relevance of this locus, we scored NL male courtship frequencies on CFS populations separated according to their genotype at this particular QTL and their respective cytoplasmic background: LL[L], LG[L], GL[G] and GG[G]. Courtship frequencies varied significantly among genotypes at this locus, with GG[G] individuals eliciting the highest frequencies and LL[L] the lowest. LG[G] individuals also induced significantly lower courtship frequencies than GG[G], whereas GL[L] individuals showed intermediate responses, not significantly different from either LG[G] or GG[G] (Fig. 5B). Interestingly, this QTL partially overlapped with a QTL for 11,x-DiMeC37 abundance, one of the key female pheromonal compounds, which also exhibited the highest LOD score at marker CHR1_2425269_1 Fig. 5C). Genotypes LL[L] and LG[G] showed significantly higher 11,x-DiMeC37 levels than GG[G] and GL[L], with no significant differences within either genotype pair (Fig. 5D). We further identified 260 genes in the overlapping QTL confidence intervals (13.3 - 36.6 cM, corresponding to 2 - 3.6 mb). Among them, *CYP4AB17* (Cytochrome P450 decarboxylase AB17) and *ACAT1* (Acetyl-CoA acetyltransferase, cytosolic) are annotated with predicted functions in CHC biosynthesis and fatty acid (FA) metabolism (Fig. 5E).

**Figure 5.**
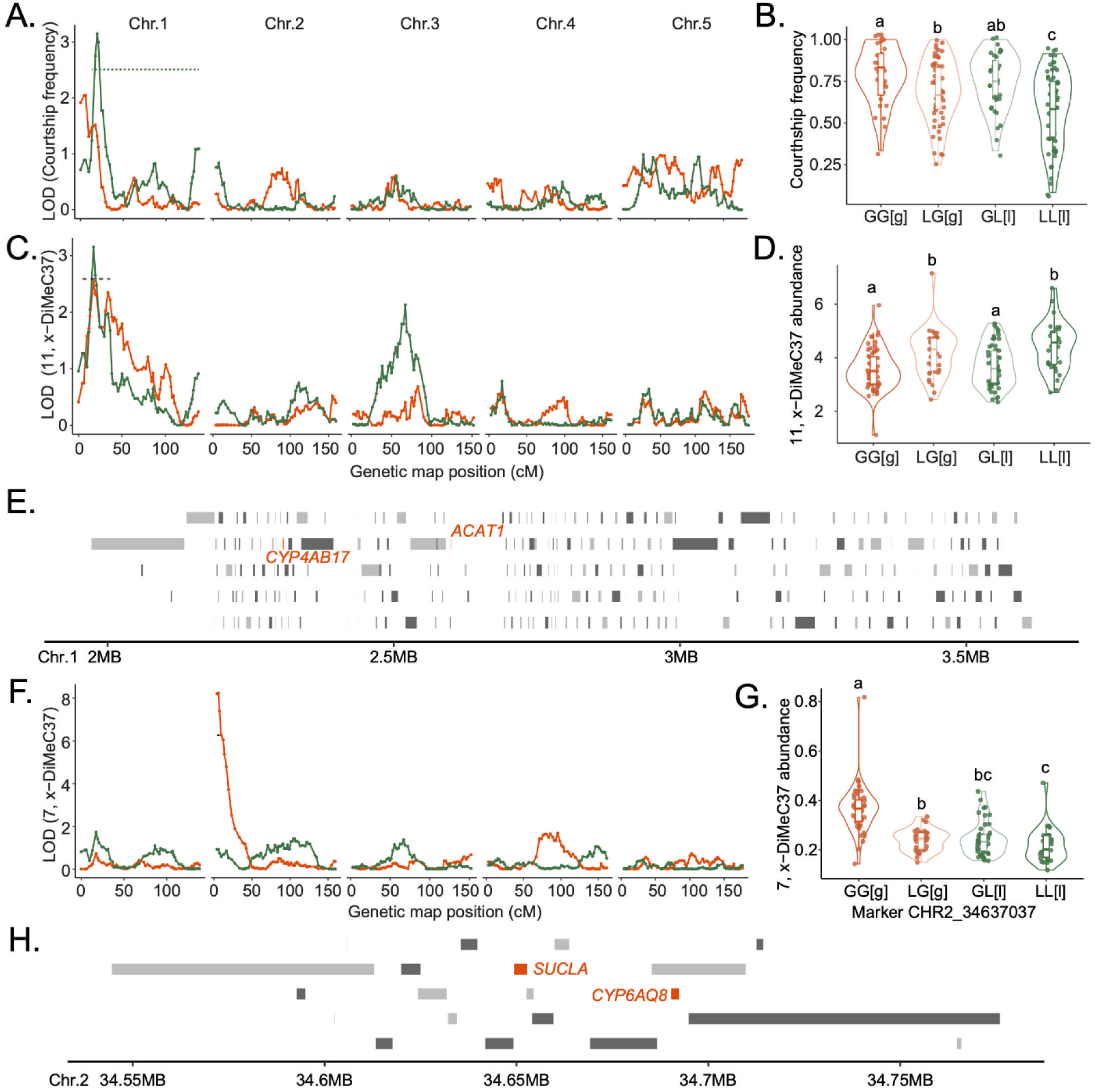
QTL localizations for male mating behavior and key pheromonal compounds in CFS mapping populations. A.) LOD (logarithm of the odds) score distribution from quantative trait loci (QTL) screening for the variation in NL male courtship frequency on CFS[L] (green) and CFS[G] (orange) populations. Positions on the genomic *Nasonia* map are indicated in centiMorgan (cM). The dotted line indicates the 95% Bayesian credible interval threshold. B.) Relative courtship frequencies of NL males on CFS populations separated according to their genotype at the detected QTL marker with the highest LOD score from A) (CHR1_2997922_1) and their respective cytoplasmic background: LL[L], LG[L], GL[G] and GG[G] indicated in orange, light orange, light green and green, respectively. Significant differences assessed with Mann-Whitney U tests and indicated by different letters. C.) LOD score distribution from QTL screening for the variation in key pheromonal compound 11,x-DiMeC37. Color codes and labels as in A). D.) Relative 11,x-DiMeC37 abundances in CFS populations separated according to their genotype at the detected QTL marker with the highest LOD score from C) (CHR1_2425269_1) and their respective cytoplasmic background, color codes, labels and significance assessment as in B). E.) Genomic region underlying the overlap between the detected QTL marker for courtship frequency (CHR1_2997922_1) and the marker for 11,x-DiMeC37 abundance (CHR1_2425269_1). Single genes are represented by grey (plus strand) and light-grey (negative strand) blocks, respectively. Candidate genes annotated for involvement in CHC biosynthesis or fatty acid metabolism (Buellesbach et al. 2022; Sun et al. 2025) are highlighted in red. F.) LOD score distribution from QTL screening for the variation in key pheromonal compound 7,x-DiMeC37. Color codes and labels as in A) and C). G.) Relative 7,x-DiMeC37 abundances in CFS populations separated according to their genotype at the detected QTL marker with the highest LOD score from F) (CHR2_34637037) and their respective cytoplasmic background, color codes, labels and significance assessment as in B) and D). H.) Genomic region underlying the detected QTL marker for 7,x-DiMeC37 abundance (CHR2_34637037). Labels and color codes for single genes as in E).

We also screened candidate genes within the QTL confidence intervals of the other two key pheromonal compounds 7,x-DiMeC37 and 13^+^-MeC37. For 7,x-DiMeC37, a QTL region was only identified in CFS[G] on chromosome 2 (3.9 to 7.3 cM; Fig. 5F). Within this region, 20 genes were identified, with *CYP6AQ8* (cytochrome P450 6AQ8) and *SUCLA* (Succinate-CoA ligase) predicted to be involved in CHC biosynthesis (Fig. 5H). GG[G] genotypes exhibited significantly higher 7,x-DiMeC37 abundance than other genotypes (Fig. 5G). No QTL was detected for 13+-MeC37 in either background.

Utilizing the generated high resolution genomic map, we also explored other candidate genes underlying genomic hotspots with co-localized QTL for structurally similar CHC compounds. On chromosome 1, QTL for the variation of 12 CHCs overlapped, primarily constituting di- and tri-methyl-branched alkanes, at a 218 kb between 54.0-56.3 cM (see Fig. 2A; Table 1). This region contained 47 genes, among which *ACADSB* (Acyl-CoA dehydrogenase for short/branched chain) and *CYP4AB6* (Cytochrome P450 decarboxylase AB6) were the most promising candidates for CHC biosynthesis (Fig. S4A). On chromosome 5, QTL for the variation of another 12 primarily methyl-branched CHC compounds overlapped at a 437 kb region between 47.3 to 58.2 cM. Among the 71 genes from this region, *fas5/6* (Fatty acid synthase), *fas6*, *fas-like* and *ACSBG1* (Long-chain-fatty-acid-CoA ligase) are the most promising CHC biosynthesis candidate genes (Fig. S4B).

### 6. QTL mapping of male CHC variation

To gain a complete picture into the genetic and phenotypic differentiation separating male and female CHC profiles, we performed a comparative QTL analysis on the CHC variation in the 126 F_2_ male fathers of the CFS lines as well (Fig. 6). CHC variation between *Nasonia* males and females has so far only been found to be quantitative (see Fig. S1), and no signaling function has hitherto been demonstrated for *Nasonia* male CHC profiles (Steiner et al. 2006; Buellesbach et al. 2013; Buellesbach 2018). We further separated the F2 male population according to their respective cytoplasmic backgrounds (*i.e.*, F2[G] and F2[L]), which, as opposed to the CFS lines (Fig. 3A), did not result in a detectable phenotypic separation based on CHC profile divergence (Fig. S5). Similarly, most QTL overlap for both backgrounds with the pooled F2 male population (Fig 6A-B). However, we also identified three QTL unique to F2[G] males and eight unique to F2[L] males, respectively.

**Figure 6.**
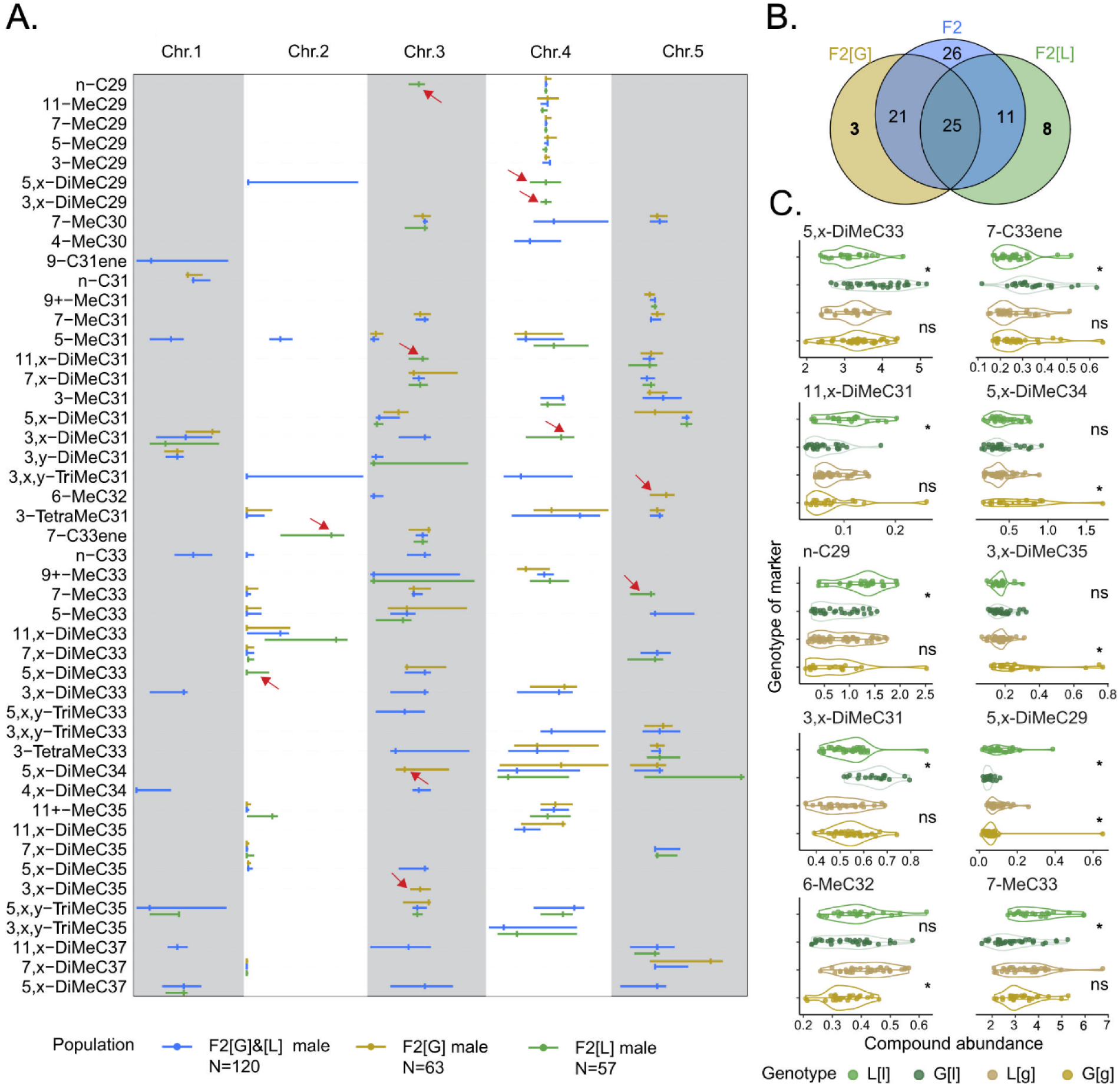
QTL mapping of CHC abundances in different F_2_ hybrid males populations. A.) QTL positions associated with CHC abundance variation depicted with 95% Bayesian credible intervals. Tick marks label the genetic marker with the highest LOD (logarithm of the odds) score within each QTL region. The general F_2_ male mapping population is indicated in blue, the F2 male population with an NG cytoplasmic background (F_2_[G]) in yellow, and the F_2_ male population with an NL cytoplasmic background in green, repectively. Color codes are consistent across panels. Red arrows indicate QTL unique in either F_2_[G] or F_2_[L] males. B.) Venn diagram of unique and shared QTL across F_2_ male hybrid mapping popuplations. C.) Variation of compound abundances for which unique QTL were identified in either F_2_[G] or F_2_[L] males, separated according to their genotype at the detected QTL marker with the highest LOD score and their respective cytoplasmic background: L[l], G[l], L[g] and G[g]. Pairwise Mann-Whitney U tests were performed to assess significant differences within matching cytoplasmic backgrounds of the mapping population genotypes: ns = not significant; * = p<0.05, N=120 for F_2_, N=63 for F_2_[G] and N=57 for F_2_[L] males.

This suggests a detectable effect of the cytoplasmic background on the genetics governing CHC variation in the haploid males just as demonstrated before in the diploid females (Fig. 3D). Compound abundances for which unique QTL with the highest LOD scores were identified in either F2[G] or F2[L] males were then further separated into their corresponding genotypes L[L], G[L], L[G] and G[G] and compared (Fig. 6C). These abundances partially varied significantly according to the respective male genotypes, revealing hitherto unrecognized cytoplasmic-haplotype interactions influencing male CHC profiles despite their lack of signaling functionality.

## Discussion

Our fine-scale investigation of female cuticular sex pheromone profiles in *Nasonia* identified three long-chained methyl-branched alkanes whose abundances significantly correlated with species-specific male mating preference. We further pinpointed genomic hotspots enriched for candidate biosynthesis genes of these key pheromonal compounds, revealing an intriguing glimpse into the potential genetic basis for species-specific chemical signaling and associated mate discrimination mechanisms.

Species-specific pheromone blends are widely used to attract and elicit mating behaviors from conspecific mates while discouraging mating with heterospecific, reproductively incompatible mating partners (Niehuis et al. 2013; Zhang et al. 2014; Xue et al. 2018). However, how biologically relevant information like species identity is encoded within complex chemical profiles remains poorly understood (Weiss et al. 2015; Wurdack et al. 2015; Würf et al. 2020). Our study bridges this knowledge gap by isolating three out of 54 total compounds in female *Nasonia* cuticular hydrocarbon (CHC) profiles as primary recognition cues for *N. longicornis* males, highly correlating with their selective mating behavior. Intriguingly, these compounds share structural features with previously identified mediators of sexual attractiveness in the closely related species *N. vitripennis* (Sun et al. 2023): all constitute long-chained methyl-branched alkanes with their first methyl group positioned at the 7th, 11th and 13th carbon atom. This suggests that similar chemical encoding mechanisms, namely positional methylation, underlie both general sexual attractiveness and species-specific recognition cues.

Quantitative differences in these compounds between females of the parental species suggest dual roles in conveying species identity and mate quality detectable by *N. longicornis* males. Interestingly, this resembles the coding strategy of moth pheromone blends, where ratios of long-chain alcohols, aldehydes, and acetate esters encode prezygotic isolation cues (Klun 1975; Tillman et al. 1999; Ando et al. 2004). Although CHC-based pheromones constitute tactile cues functioning at close range while moth pheromones are long-range volatiles, the genetic and biosynthetic origins of these compound classes bear striking similarities (Roelofs et al. 2002; Lienard et al. 2010; Hagström et al. 2013). For instance, fatty-acyl CoA reductases (FARs) are essential in the biosynthesis of long-chain fatty alcohols in moths (Lienard et al. 2010) as well as in converting fatty acyl-CoAs into primary alcohols as an intermediate step in CHC biosynthesis (Holze et al. 2021). Exploring further genetic and biosynthetic parallels between these evolutionarily distant systems separated by more than 330 million years could uncover deeply conserved elements of insect chemical communication (Misof et al. 2014).

Correlation networks centered on the three key pheromonal compounds showed stronger sex- than species-specific differences. This aligns with prior findings and reflects the sex-limited nature of these chemical profiles: only female CHCs function in sexual communication, likely resulting in distinct selective pressures separating the sexes (Steiner et al. 2006; Buellesbach et al. 2013; Buellesbach et al. 2018b). Concordantly, the identified pheromonal compounds showed strong correlations in CFS females which were generally absent or weaker in F2 males. In addition, we uncovered effects of allelic interactions (*i.e.,* dominant/recessive effects) and cytoplasmic background on CHC variations unrecognized in previous studies (Niehuis et al. 2011; Buellesbach et al. 2022). Among 88 QTL detected in total, only 14 were shared between CFS[G] and CFS[L], all governing methyl-branched alkane variation. Notably, for the pheromonal compound with a simultaneous impact on male courtship and copulation behavior, 7,x-DiMeC37, a QTL was only detected in CFS[G] females. Compound abundances were significantly higher in individuals with the GG genotype, indicating a strong recessive effect of the NG allele. In contrast, no effect of the cytoplasmic/haplotypic background impacting 7,x-DiMeC37 abundances could be found. However, different cytoplasmic/haplotypic backgrounds prominently affected the abundance of another key pheromonal compound, 11,x-DiMeC37. This illustrates the complexity of reliably predicting and assessing the potential impact of different genetic factors on the phenotypic traits under investigation.

We expected female sexual attractiveness in clonal female sibship lines (CFS[L] and CFS[G]) to fall between the parental *N. giraulti* and *N. longicornis* females, consistent with their intermediate CHC profiles. Surprisingly, despite clustering closer to *N. giraulti* female CHC profiles, CFS[G] lines elicited higher mating frequencies from *N. longicornis* males than CFS[L] lines. While this initially seems counterintuitive, this pattern may indicate that the investigated traits lie outside the range of the respective parental species (Kristensen and Sørensen 2005; Kaeppler 2012). In this case, these traits can either be detrimental and underperform in comparison to the parental species (*i.e.,* phenotypic disruption), or they can outperform the latter, most commonly resulting in higher reproductive outputs (*i.e.,* heterosis/’hybrid vigor’) (Fitzpatrick 2008; Ekechukwu et al. 2015; Fezza et al. 2022). These effects are frequently observed in various traits of hybrids between various parasitoid species (Legner 1972; Hoy 1975; Benvenuto et al. 2012; Tsutsui et al. 2014). If this is also the case here, it appears likely that either the CFS[G] lines display some degree of hybrid vigor, or the CFS[L] lines some degree of phenotypic disruption, or a combination of both. Additional asymmetry arose from diapause induction: a large proportion of offspring from backcrosses to NL females entered prolonged diapause as opposed to the backcrosses to NG females. This offspring did not further develop into adults and did therefore also not contribute to the generation of CFS[L] lines. Therefore, larger numbers of CFS[L] populations needed to be generated to achieve equal proportions for both cytoplasmic backgrounds in our experimental lines. This not only indicates stronger postzygotic isolation in the direction of NL, as demonstrated before (Breeuwer and Werren 1995; Ellison et al. 2008; Gibson et al. 2013), but also an additional bottleneck impacting hybridization dynamics and potentially lowering variation in our CFS[L] lines.

Furthermore, a QTL indirectly influencing *N. longicornis* male courtship frequencies was detected only in the CFS[L] population. This was driven by large differences in elicited male courtship frequencies between LG[L] and LL[L] genotypes, whereas the analogous genotypes in CFS[G] (GG[G] vs. GL[G]) exhibited much smaller effects, below the threshold of QTL detection. Moreover, the correlation networks between the three key pheromonal compounds and other CHCs differed markedly between CFS[G] and CFS[L], suggesting diverging regulatory governance of CHC quantities between these cytoplasmic and haplotypic backgrounds. This highlights how hybrid phenotypes may be shaped by complex, non-additive interactions, likely reflecting unresolved incompatibilities between diverged parental genomes.

Within QTL regions associated with the three uncovered key pheromonal compounds, we shortlisted several potential CHC biosynthesis candidate genes. Most notably, the narrow QTL region governing 7,x-DiMeC37 abundance contains *CYP6AQ8* (a cytochrome P450 decarbonylase) and *SUCLA* (succinate-CoA ligase). The cytochrome P450 decarbonylase gene subfamily constitutes a vital component of the CHC biosynthesis pathway, catalyzing the oxidative decarbonylation of CHC precursors (Feyereisen 2012; Qiu et al. 2012; Holze et al. 2021). Succinate-CoA ligases, on the other hand, catalyze the reversible conversion of succinyl-CoA to succinate, which plays a critical role in the citric acid cycle (Johnson et al. 1998), whose key substrate acetyl-CoA serves as the main and rate-limiting precursor for CHC biosynthesis (Barber et al. 2005; Holze et al. 2021). Similarly, the QTL region impacting both 11,x-DiMeC37 abundance and *N. longicornis* male courtship frequency contains another P450 enzyme (*CYP4AB17*) and *ACAT1*, which regulates acetyl-CoA pools via acetoacetyl-CoA formation.

Intriguingly, a QTL region on chromosome 5 shared across both CFS backgrounds contains orthologues of the tandem duplicated fatty acid synthase genes *fas5/6* from *N. vitripennis*. Knockdown of *fas5/6* in this closely related *Nasonia* species caused pronounced shifts in female CHC profiles specifically altering the ratios of methyl-branched CHCs similar to those identified in the present study (Sun et al. 2023). Notably, this knockdown also led to a marked reduction in female sexual attractiveness to conspecific males. Taken together, these findings suggest conserved molecular pathways underlying both intraspecific sexual attractiveness and interspecific mate discrimination.

Comparing the QTL governing CHC variations between F_2_ haploid fathers and their F_3_ diploid CFS daughters reveal further striking differences: more than half of the QTL detected in females are unique and did not overlap with any QTL detected in the males. Such pronounced differences in QTL numbers and mapping locations were unexpected and surprising, particularly in a system where sexes are determined by ploidy level(Henter 2003; Tien et al. 2015). Largely transcending mere allelic effects, these findings highlight the need to strictly separate male and female genetics rather than extrapolating across sexes in similar future genomic studies.

In conclusion, our study advances our understanding of the genetic and chemical basis of species-specific pheromone blends and associated mate discrimination mechanisms. The importance of sex-specific regulatory effects, cytoplasmic backgrounds, and genomic interactions that also emerged from our study highlight the need for careful experimental designs that account for these layers of complexity in future studies. In addition, our findings provide novel insights into how biological information is encoded in complex chemical profiles, and how subtle divergence in signaling molecules can reinforce species boundaries and might have contributed to early speciation processes.

## Materials and methods

### 1. *Nasonia* strain maintenance

The standard *Wolbachia*-cured laboratory strains RV2(u) and IV7(u) that represent the species *Nasonia giraulti* (NG) and *N. longicornis* (NL), respectively, were used in this study. Strains were maintained at 25 degrees under a 16:8h light-dark cycle. *Calliphora vomitoria* pupae (Diptera: Calliphoridae) served as hosts for wasp rearing.

### 2. Hybrid crossing scheme

Reciprocal interspecific crosses between NG and NL were performed as described in Buellesbach et al. (2022). Virgin males and females were isolated at the black pupal stage and housed individually until emergence. Within 0-24 h after emergence, reciprocal crosses (NG male x NL female; NL male x NG female) were set up, and after further 24 h for mating, males were removed and host pupae were provided for females to oviposit. F_1_ virgin females were collected after 12 days (black pupal stage), isolated and allowed to lay unfertilized eggs after emergence, producing haploid F_2_ hybrid males. Each thus generated F_2_ male was backcrossed sequentially to a respective female from both parental strains (NG and NL), allowing a total of 24 h for mating per female. The mated females were removed and provided with six host pupae. Consequently, F_3_ female hybrid offspring from each of the mated F_2_ females collectively constituted a single clonal female sibship line, which were collected at the black pupal stage, freeze-killed, and stored at −80 °C for further analysis. Similarly, their haploid F_2_ male hybrid fathers were collected, freeze-killed, and stored at −80 °C. Freeze-killing was performed with liquid nitrogen.

### 3. Genotyping of F_2_ hybrid males

Restriction-site associated DNA (RAD) sequencing was performed on the F_2_ hybrid male fathers of the CFS lines. DNA was extracted using the Qiagen DNeasy Blood & Tissue Kit (Qiagen, Hilden, Germany). Genomic libraries were constructed following Brelsford et al. (2016). Briefly, genomic DNA was digested with the two restriction enzymes EcoRI and MseI ligated to adaptors with sample-specific barcodes, and amplified via PCR using Q5 Hot Start Polymerase® (New England Biolabs GmbH Frankfurt, Germany) for 20 cycles. Primers and dNTPs were added to the final thermal cycle to reduce single-stranded or heteroduplex PCR products. After total DNA volume reached 7 µl per reaction, the PCR products were pooled, and fragments insert sizes ranging between 300-700 bp (including 128 bp of adaptors and primers) were selected using BluePippin (Sage Science, Inc., Beverly, MA, USA). Libraries were sequenced with paired-end 150 bp reads on an Illumina HiSeq X platform. Raw sequence quality was assessed using FastQC (v.0.11.7). Demultiplexing was performed with the command *process_radtags* from the Stacks2 pipeline(Rochette et al. 2019). The resulting reads were mapped to the *Nasonia vitrpennis* genome assembly *Nvit_psr_1.1* (GenBank assembly accession: GCA_009193385.2), using Bowtie2 (Langmead and Salzberg 2012). Reads with multiple hits in the reference genome were filtered out. SNP calling from the mapped sequences was performed using the command *ref_map.pl* from the Stacks2 pipeline (v2.62) and genotype data was subsequently filtered using vcftools (v0.1.13) (Danecek et al. 2011) with the following parameters: maf - 0.35; maxDP - 16; minDP - 2; max-missing - 0.8. Genotypes of inbred NG and NL males served as references for parental allele assignment. Detailed pipelines are available in the associated Figshare repository.

### 4. Chemical analysis

Cuticular hydrocarbons (CHCs) were extracted by immersing individual wasps in 50 µl HPLC-grade *n*-hexane (Merck, KGaA, Darmstadt, Germany) in 2 ml GC-MS glass vials (Agilent Technologies, Waldbronn, Germany) on an orbital shaker (IKA KS 130 Basic, Staufen, Germany) for 10 minutes. Subsequently, extracts were evaporated under a continuous stream of gaseous carbon dioxide and then reconstituted in 10 μL of a hexane solution containing 7,5 ng/ μL dodecane (C12) as an internal standard. Following this, 3 μL of the reconstituted extract was injected in splitless mode with an automatic liquid sampler (PAL RSI 120, CTC Analytics AG, Zwingen, Switzerland) into a gas-chromatograph (GC: 7890B) simultaneously coupled to a flame ionization detector (FID: G3440B) and a tandem mass spectrometer (MS/MS: 7010B, all provided by Agilent Technologies, Waldbronn, Germany). The system employed a fused silica column (DB-5MS ultra inert; 30 m x 250 μm x 0.25 μm; Agilent J&W GC columns, Santa Clara, CA, USA) with helium serving as the carrier gas under a constant flow of 1.8 ml/min. The FID operated at 300 °C and used nitrogen as make-up gas (20 mL/min), and hydrogen as fuel gas (30 mL/min). The column was split at an auxiliary electronic pressure control (Aux EPC) module into two deactivated fused silica column pieces (0.9 m x 150 μm and 1.33 m x 150 μm) with flow rates of 0.8 mL/min and 1.33 mL/min, respectively, leading into the FID detector and the mass spectrometer. The column temperature program started at 60 °C and held for 1 min, then increased at a rate of 40 °C per minute until 200 °C, followed by a gradual increase of 5 °C per minute to a temperature of 320 °C, held for 5 min.

CHC peak detection, integration, quantification, and identification were all carried out with Quantitative Analysis MassHunter Workstation Software (Version B.11.00, Agilent Technologies, Santa Clara, California, USA). CHCs were identified according to their retention indices, diagnostic ions, and mass spectra derived from the total ion count (TIC) chromatograms, whereas quantifications used FID chromatograms. Absolute CHC quantities (in ng) were obtained by calibrating each compound according to a dilution series based on the closest eluting *n*-alkane from a C21-40 standard series (Merck, KGaA, Darmstadt, Germany) at 0.5, 1, 2, 5, 10, 20, 40 ng/ μL, respectively. The relative abundance of single compounds was calculated by dividing their absolute quantity to overall CHC quantity in each sample.

### 5. Behavioral assays

Sexual attractiveness of females was tested by assessing courtship (head nods and antennal sweeps) and copulation frequencies on freeze-killed female dummies, which have been established as main indicators of male mate acceptance and female sexual attractiveness (Buellesbach et al. 2013; Buellesbach et al. 2018b; Sun et al. 2023). NL males have previously demonstrated discriminatory behavior between con- and heterospecific (NG) females (Giesbers et al. 2013). To confirm this behavior, an initial discriminatory experiment was set up pairing virgin NL males with 119 NL and 120 NG females, respectively, and scoring courtship and copulation. After confirmation, NL males were employed for assessing sexual attractiveness of each clonal female sibship (CFS) population, assaying twelve females per population. Assays were conducted in mating chambers comprising two aluminum plates (53 × 41 × 5 mm), each with 12 observation holes (6 mm diameter) that served as observation sites. Individual freeze-killed female dummies were randomized and placed in each hole of one of the two plates, virgin NL males in each hole of the opposite plate and then immediately covered with glass slides (Diagonal GmbH &Co. KG, Münster, Germany). Behavioral assays were initiated by quickly adjoining the two plates, and then record the male behavior on the female dummies for 5 mins with a Canon EOS 70D camera (Canon inc., Tokyo, Japan). All behavioral assays were performed in an enclosed wooden box with constant illumination (100Lm, LED Dioder light L0601, IKEA, Delft, The Netherlands). Significant differences in courtship and copulation behavior were assessed with Fisher’s exact tests (NL males on NL vs. NG females), correlations of courtship and copulation displays with individual CHC compound abundances were assessed with Benjamini-Hochberg corrected Spearman rank correlation tests (NL males vs. CFS populations).

### 6. Correlation network analysis

We further investigated the phenotypic interactions between the detected key pheromonal compounds and other CHCs in the CFS populations as well as their F_2_ hybrid male fathers. Benjamini-Hochberg-corrected Spearman rank correlation tests were performed between the relative abundance of all 54 CHC compounds. Networks between the three key pheromonal compounds and other connected CHCs were visualized using the R package *igraph* (version 0.10.10) (Csardi and Nepusz 2005). Only the correlations with an adjusted Spearman correlation coefficient larger than 0.5 or smaller than -0.5 were visualized as edges in the networks.

### 7. Quantitative trait loci mapping and candidate gene identification

Quantitative trait loci (QTL) mapping for CHC variation and male mating behavior was performed using R/qtl2 (Broman et al. 2019). Haley-Knott regression (Haley and Knott 1992) was applied to each marker in the genetic map with an assumed genotype error probability of 0.002 and a fixed step width of 1 cM. Statistical significance of QTL presence for each compound was estimated and adjusted from 1000 permutations. Bayesian credible interval (95%) were calculated for approximate locations of every detected QTL. Furthermore, we assessed the percentage of phenotypic variance explained (PVE) by the QTL using the following formula:

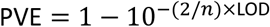

where *n* corresponds to the sample size of the mapping individuals and LOD to the logarithm of the odds score. QTL mapping was performed separately for CFS[G] and CFS[L] as well as their F_2_ hybrid male fathers, using genotypic data and genetic maps acquired from the latter. Physical positions of QTL were aligned to the *N. vitripennis* Nvit 2.0 reference genome (Rago et al. 2016), and selected candidate genes annotated for CHC biosynthesis or fatty acid metabolism (Buellesbach et al. 2022; Sun et al. 2025) were highlighted.

## Author Contributions

J.B. & J.G. conceived the study; J.B. & W.S. designed the study; W.S., M.F. & A.T performed the experiments and collected the data, W.S. analyzed the results; W.S. validated the results; W.S. visualized the data; J.B. & J.G. provided the resources; W.S. & J.B. wrote the original draft; J.B., W.S., E.P. & J.G. reviewed and edited the final manuscript version, J.B. & J.G. supervised the project; J.B. acquired the funding for the study.

## Acknowledgement

We acknowledge Kathrin Brüggemann for the chemical extraction of thousands of samples. We also appreciate Mohammed "Simo" Errbii for the help and accompany in the bioinformatic analysis. Additionally, we thank Marek Golian and Ming Zhou for the sample collection during the construction of hybrid populations.

## Supplementary information

**Figure S1.**
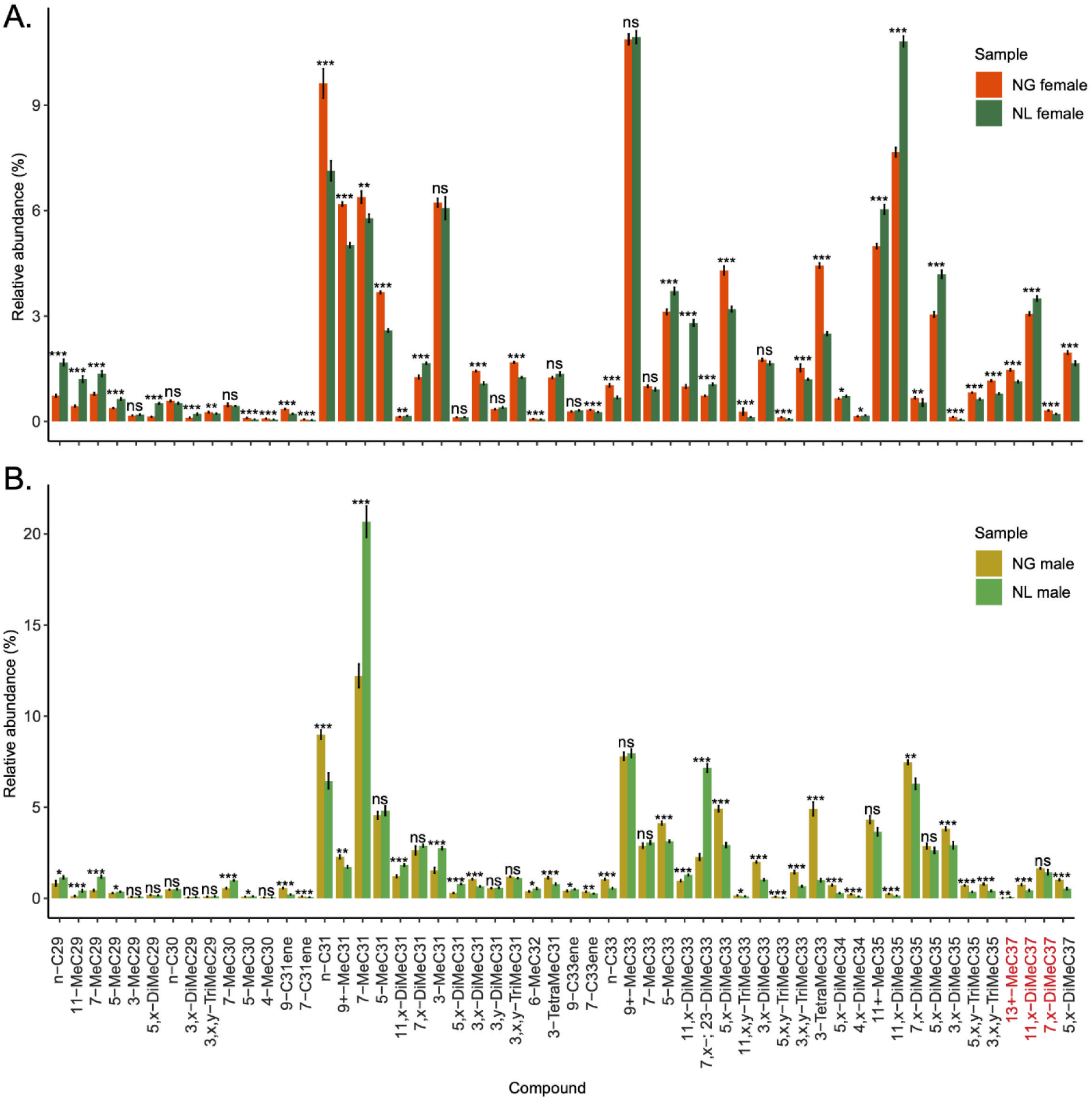
Relative abundances of identified CHC compounds occurring in NL and NG females (A) and males (B). Abundance calculated as proportion of total CHC quantity. Benjamini-Hochberg corrected Mann-Whitney U tests were used to assess significant differences, indicated as * for p<0.05; ** for p<0.01; *** for p<0.001; ns for not significant. Females: NG (red), NL (dark green); males: NG (yellow), NL (light green); N = 10 per group.

**Figure S2.**
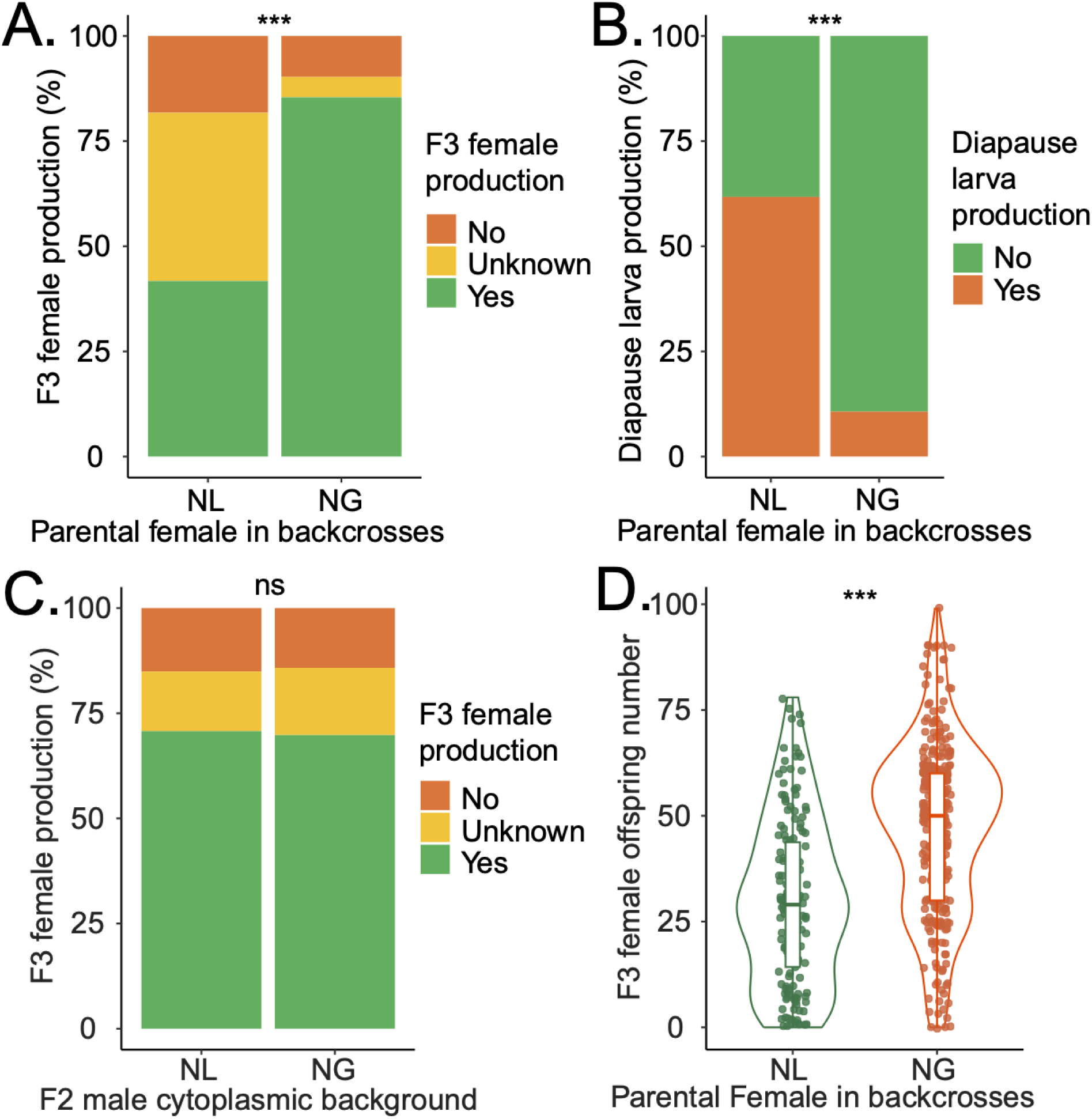
Cytoplasmic incompatibility in F_3_ clonal female sibship (CFS) populations. A.) Comparison of relative F_3_ CFS offspring production from NL and NG females. ‘Unknown’ refers to the percentage of diapause larvae (see B) that do not further develop into differentiable adults. B.) Proportion of diapause larvae produced by NL and NG females C.) Effect of F_2_ male cytoplasmic background (F_2_[L] or F2[G]) on CFS offspring production. D.) Female offspring numbers in each CFS population produced by NL and NG females. Statistical significance was assessed by Fisher exact tests for A.), B.) and C.), and a Mann-Whitney U test in D.) Significance levels are indicated as follows: * for p<0.05; ** for p<0.01; *** for p<0.001; ns for not significant.

**Figure S3.**
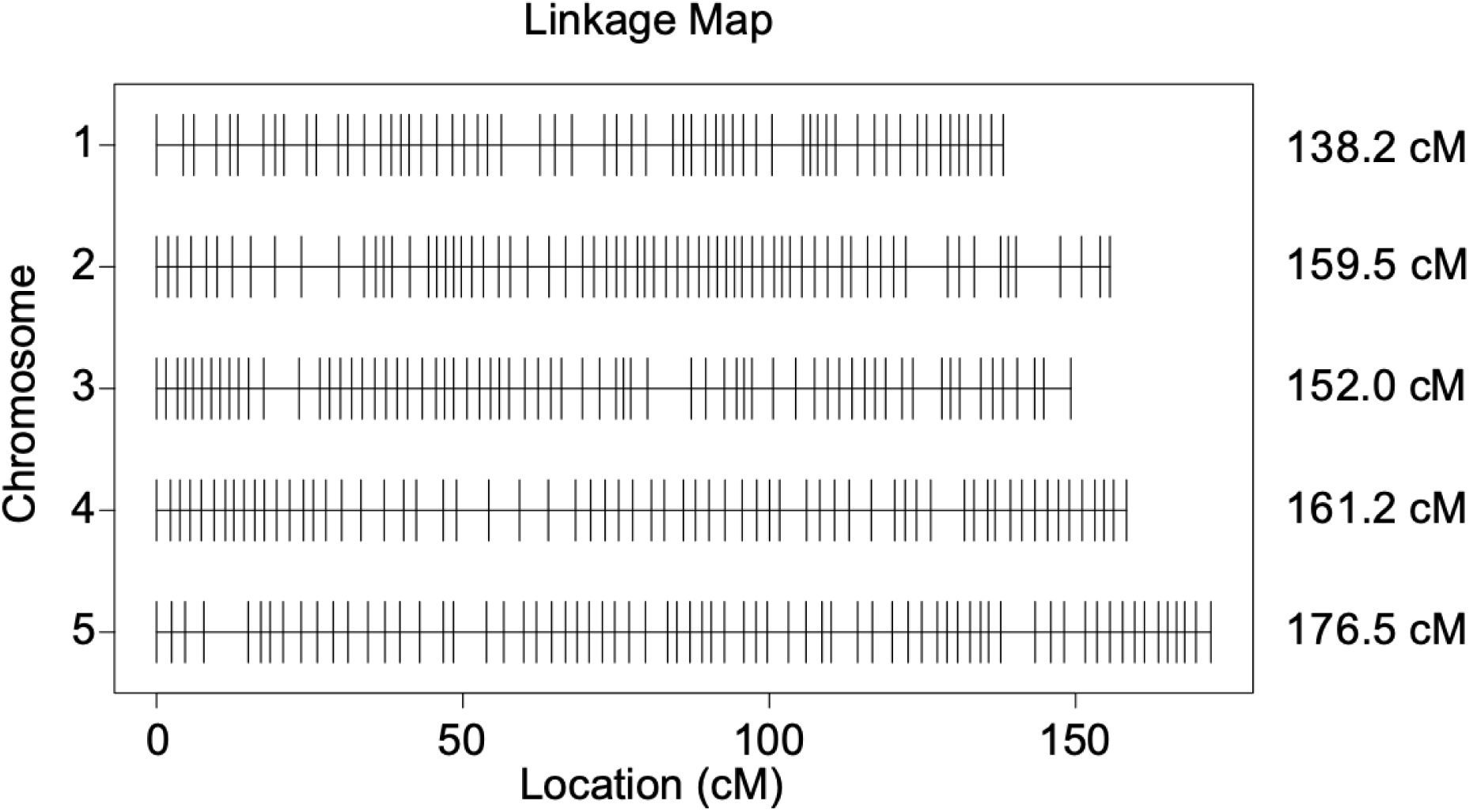
Linkage map produced from 332 SNP markers in the F_2_ male hybrid population. The genetic map consists of five linkage groups, representing the five *Nasonia* chromosomes. The length of each linkage group is indicated in centiMorgan (cM), non-recombining SNP markers are indicated as vertical lines.

**Figure S4.**
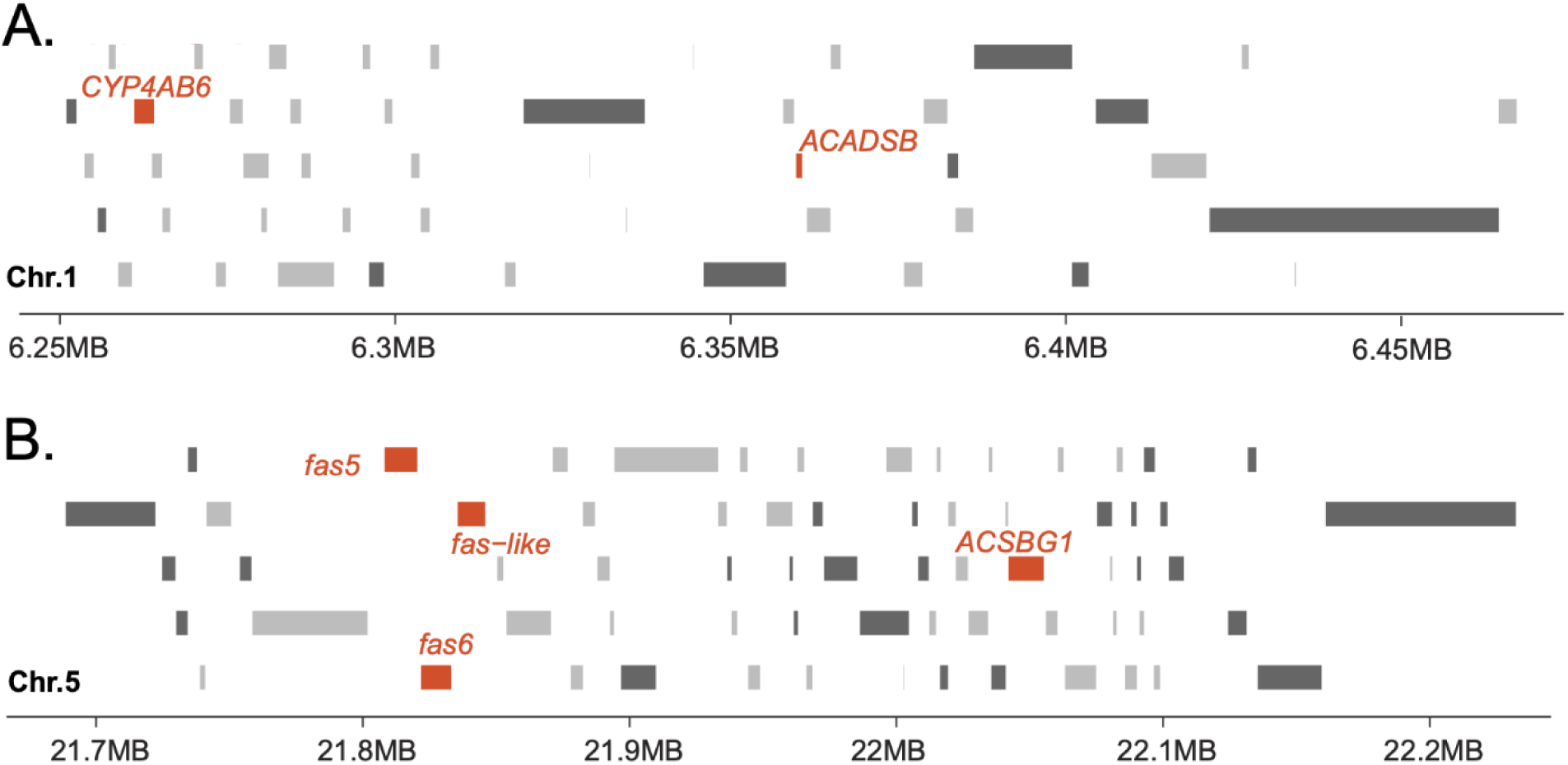
CHC biosynthesis candidate genes in overlapping QTL regions for species-specific CHC variation. A.) Genomic region on chromosome 1 between 54.0-56.3 cM (compare to Fig. 2) (6.25 to 6.45 MB) harboring co-localized QTL for the variation of 12 structurally similar, primarily di- and tri-methyl-branched CHCs. B.) Genomic region on chromosome 5 between 47.3 to 58.2 cM (compare to Fig. 2) (21.7 to 22.2 MB) harboring co-localized QTL for the variation of 12 structurally similar, primarily methyl-branched CHCs. Single genes are represented by grey (plus strand) and light-grey (negative strand) blocks, respectively. Candidate genes annotated for involvement in CHC biosynthesis or fatty acid metabolism (Buellesbach et al. 2022; Sun et al. 2025) are highlighted in red.

**Figure S5.**
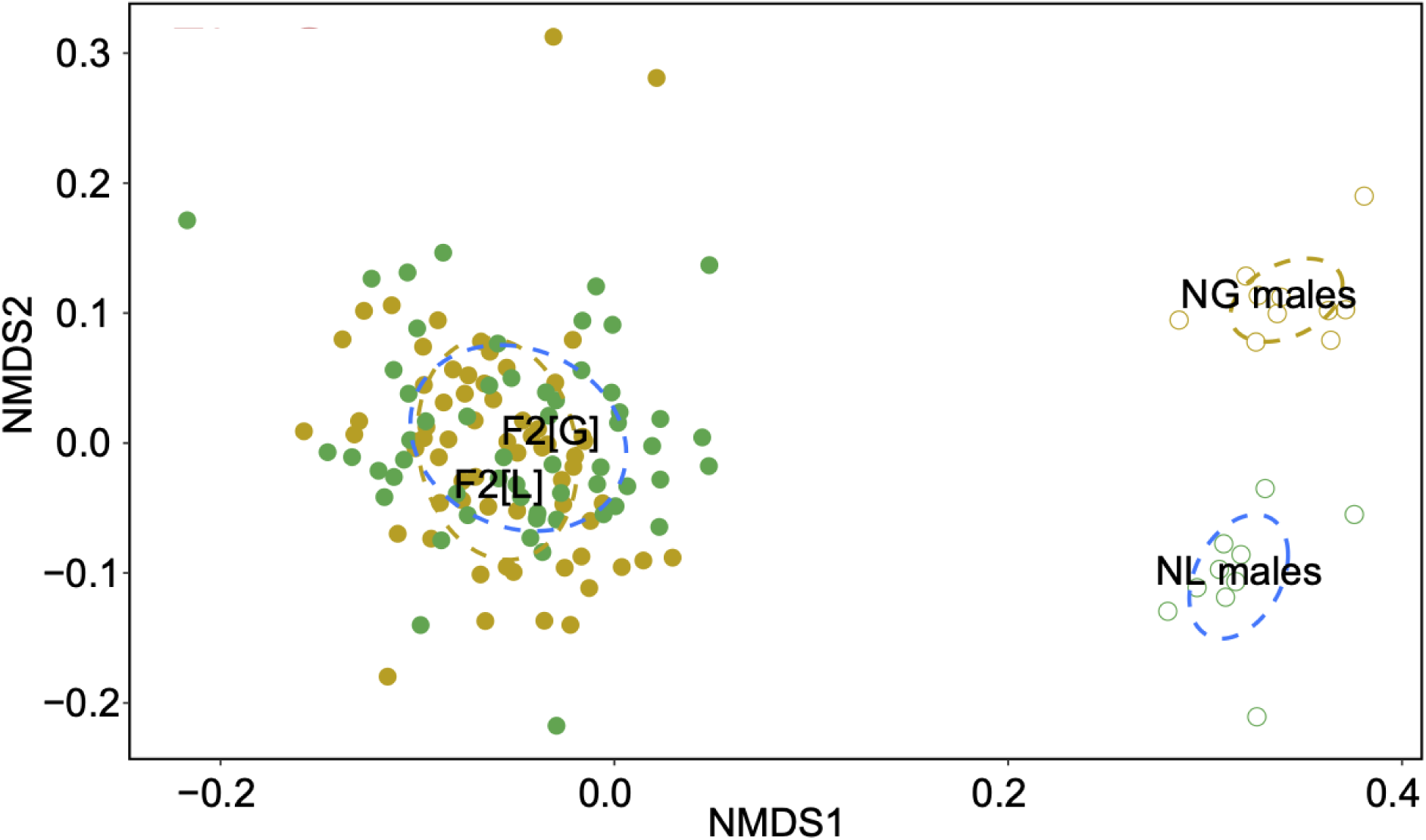
CHC profile divergence between male populations. Non-Metric Multidimensional Scaling (NMDS) plot demonstrating the divergence of CHC profiles of NL (green open circles, N=10), NG (yellow open circles, N=10), F_2_[L] (green filled circles, N=57) and F_2_[G] (yellow filled circles, N=63) male populations. Each data point represents the CHC profile of a single individual. Bray–Curtis dissimilarity model, stress value = 0.15.

**Table S1.**
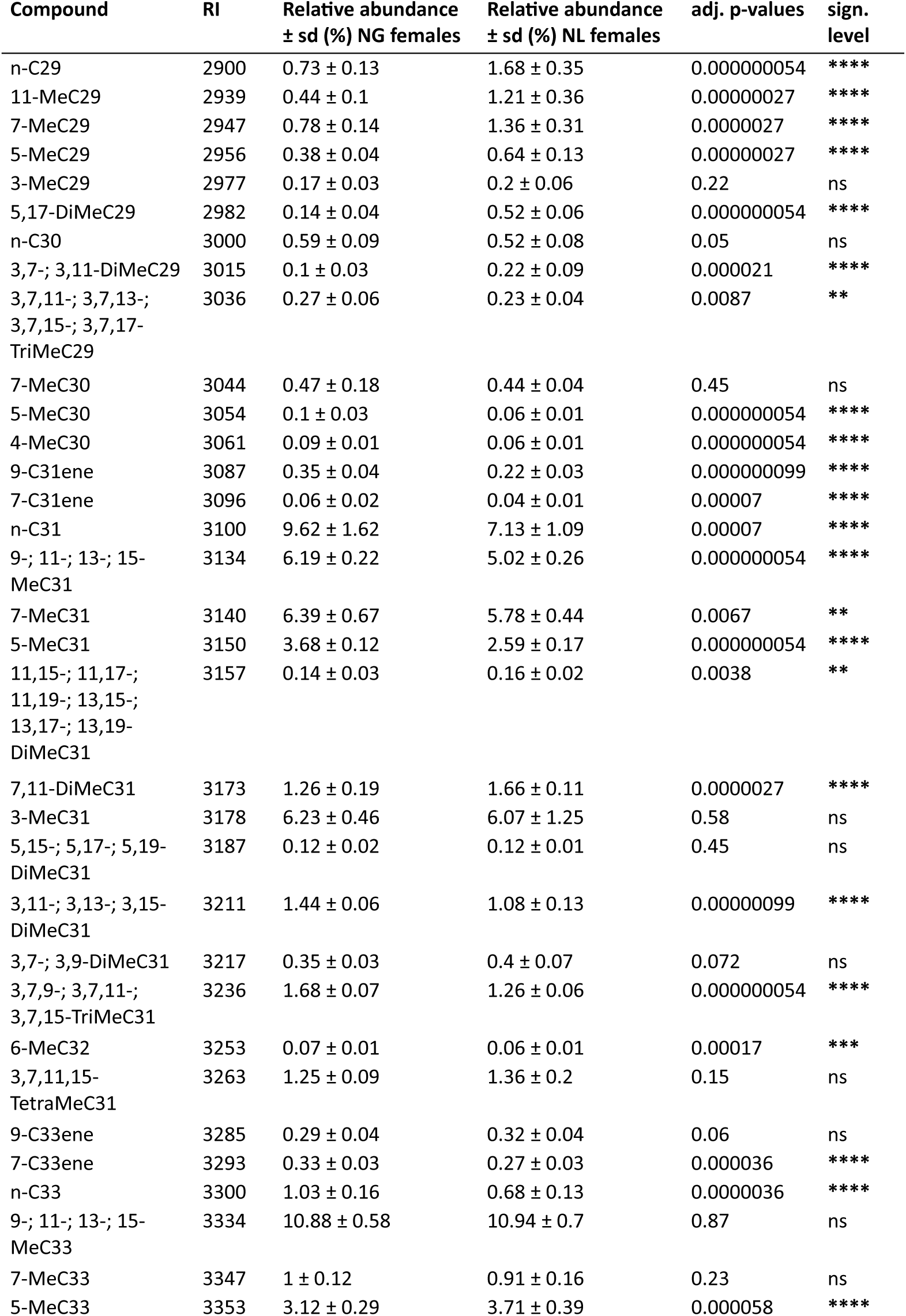
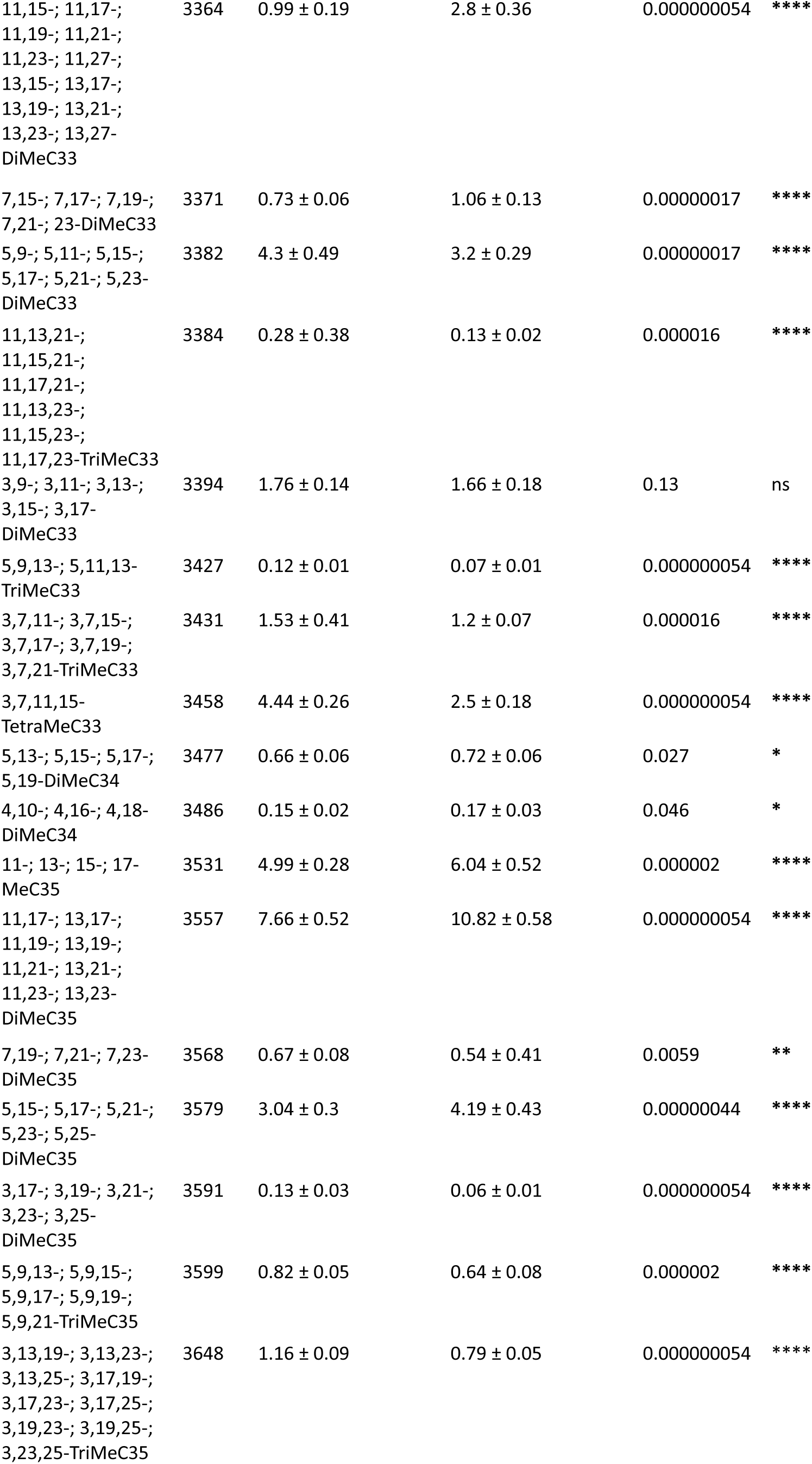
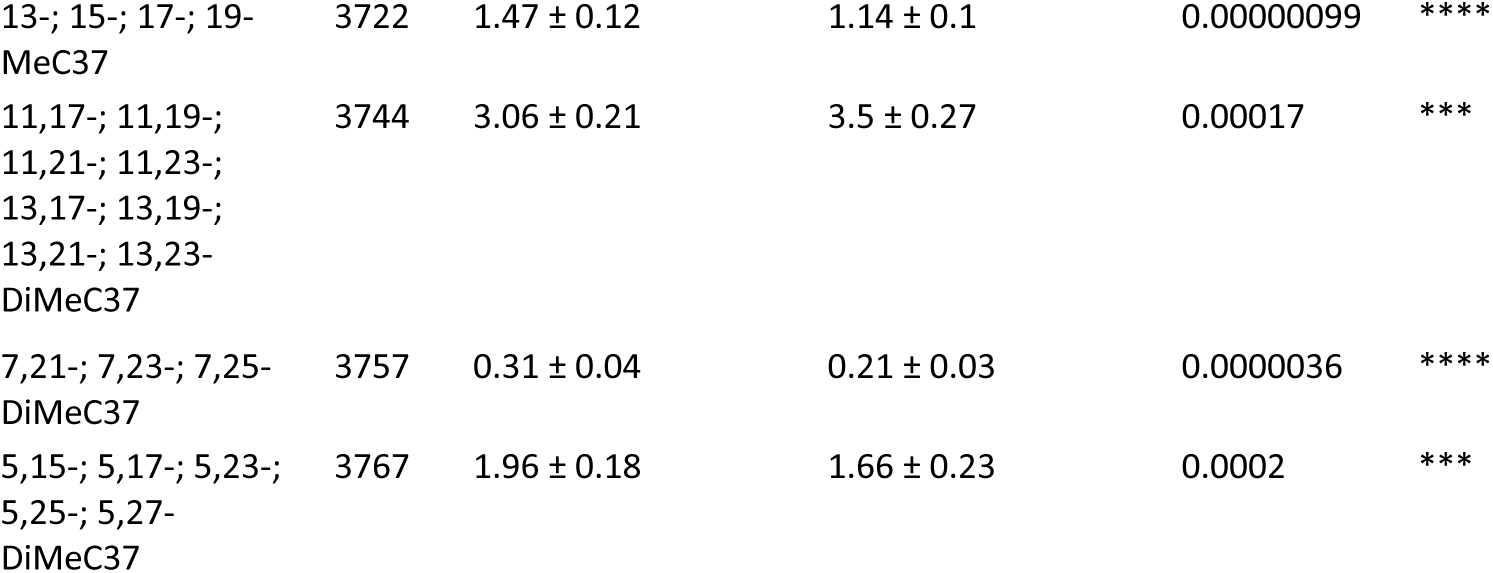
Relative abundances of CHC compounds in *N. longicornis* and *N. giraulti* females. Compound names, retention indexes (RI), mean relative abundances with standard deviations for each species, adjusted (adj.) p-values and significance (sign.) levels are indicated. Statistical significance between absolute compound abundances in *N. longicornis* and *N. giraulti* females assessed through Benjamini-Hochberg corrected Mann-Whitney U tests.

**Table S2.** Correlation statistics between CHC compound abundances in CFS lines and NG male courtship frequency. Correlation coefficients (R) and unadjusted p-values as well as corrected p-values based on Spearman rank correlation tests for each CHC compound are indicated. Corrected p-values were adjusted for multiple testing through the Benjamini-Hochberg method. Compounds showing significant correlations after p-value adjustment are highlighted in bold.

| Compound | R | p-value | corr. p-value |
| --- | --- | --- | --- |
| 11-; 13-; 15-; 17-MeC35 | 0.0068 | 0.9239 | 0.9413 |
| 11-DiMeC31 | -0.0952 | 0.1776 | 0.3995 |
| 11-DiMeC33 | -0.1726 | 0.0140 | 0.0841 |
| 11-DiMeC35 | -0.0805 | 0.2549 | 0.5294 |
| **11-DiMeC37** | **0.2119** | **0.0025** | **0.0445** |
| 11-MeC29 | -0.0957 | 0.1757 | 0.3995 |
| 11-TriMeC33 | -0.0495 | 0.4843 | 0.7069 |
| **13-; 15-; 17-; 19-MeC37** | **0.2228** | **0.0014** | **0.0387** |
| 3-DiMeC29 | -0.0317 | 0.6542 | 0.7769 |
| 3-DiMeC33 | 0.1368 | 0.0522 | 0.2013 |
| 3-DiMeC35 | 0.1837 | 0.0089 | 0.0712 |
| 3-MeC29 | -0.0389 | 0.5825 | 0.7672 |
| 3-MeC31 | -0.1000 | 0.1568 | 0.3967 |
| 3-TriMeC29 | 0.0638 | 0.3673 | 0.6175 |
| 3-TriMeC31 | 0.1796 | 0.0106 | 0.0712 |
| 3-TriMeC33 | 0.1233 | 0.0804 | 0.2714 |
| 3-TriMeC35 | 0.1814 | 0.0098 | 0.0712 |
| 3,11-DiMeC31 | 0.1317 | 0.0617 | 0.2222 |
| 3,7,11,15-TetraMeC31 | -0.1388 | 0.0489 | 0.2013 |
| 3,7,11,15-TetraMeC33 | 0.1191 | 0.0913 | 0.2738 |
| 3,9-DiMeC31 | -0.0160 | 0.8210 | 0.8526 |
| 4-DiMeC34 | 0.0517 | 0.4653 | 0.6980 |
| 4-MeC30 | -0.0310 | 0.6618 | 0.7769 |
| 5-DiMeC29 | -0.1581 | 0.0247 | 0.1332 |
| 5-DiMeC31 | 0.1204 | 0.0878 | 0.2738 |
| 5-DiMeC33 | 0.1125 | 0.1110 | 0.3155 |
| 5-DiMeC34 | -0.0252 | 0.7222 | 0.7959 |
| 5-DiMeC35 | -0.0280 | 0.6924 | 0.7956 |
| 5-DiMeC37 | 0.1866 | 0.0078 | 0.0712 |
| 5-MeC29 | -0.0989 | 0.1616 | 0.3967 |
| 5-MeC30 | -0.0836 | 0.2366 | 0.5111 |
| 5-MeC31 | -0.0014 | 0.9839 | 0.9839 |
| 5-MeC33 | -0.0365 | 0.6056 | 0.7769 |
| 5-TriMeC33 | 0.0704 | 0.3194 | 0.6161 |
| 5-TriMeC35 | 0.2012 | 0.0041 | 0.0552 |
| 6-MeC32 | 0.0464 | 0.5124 | 0.7215 |
| 7-C31ene | -0.0610 | 0.3888 | 0.6175 |
| 7-C33ene | -0.0165 | 0.8157 | 0.8526 |
| 7-DiMeC31 | -0.0639 | 0.3665 | 0.6175 |
| 7-DiMeC33 | -0.0337 | 0.6342 | 0.7769 |
| 7-DiMeC35 | 0.1421 | 0.0436 | 0.2013 |
| **7-DiMeC37** | **0.2884** | **0.0000** | **0.0017** |
| 7-MeC29 | -0.0539 | 0.4463 | 0.6886 |
| 7-MeC30 | -0.0654 | 0.3548 | 0.6175 |
| 7-MeC31 | 0.0212 | 0.7642 | 0.8253 |
| 7-MeC33 | 0.0394 | 0.5780 | 0.7672 |
| 9-; 11-; 13-; 15-MeC31 | -0.0342 | 0.6294 | 0.7769 |
| 9-; 11-; 13-; 15-MeC33 | 0.0772 | 0.2748 | 0.5496 |
| 9-C31ene | -0.0624 | 0.3775 | 0.6175 |
| 9-C33ene | -0.1396 | 0.0475 | 0.2013 |
| n-C29 | -0.1033 | 0.1435 | 0.3873 |
| n-C30 | -0.0620 | 0.3810 | 0.6175 |
| n-C31 | -0.0454 | 0.5211 | 0.7215 |
| n-C33 | -0.0265 | 0.7079 | 0.7959 |

